# Resources for molecular studies of unculturable obligate biotrophic fungal plant pathogens using their saprotrophic relatives

**DOI:** 10.1101/2025.05.08.652889

**Authors:** Anne Loos, Ella Doykova, Jiangzhao Qian, Florian Kümmel, Heba Ibrahim, Levente Kiss, Ralph Panstruga, Stefan Kusch

**Affiliations:** Unit of Plant Molecular Cell Biology, Institute for Biology I, RWTH Aachen University, Worringerweg 1, D-52056 Aachen, Germany; current address: Department of Plant-Microbe Interactions, Max Planck Institute for Plant Breeding Research, Carl-von-Linné-Weg 10, Cologne D-50829, Germany; Institute of Bio- and Geosciences IBG-2, Forschungszentrum Jülich, D-52425 Jülich, Germany; Centre for Crop Health, Institute for Life Sciences and the Environment, University of Southern Queensland, Toowoomba, QLD, Australia; Plant Protection Institute, Centre for Agricultural Research, Eötvös Loránd Research Network, Budapest, Hungary; current address: Institute of Bio- and Geosciences IBG-4, Forschungszentrum Jülich, D-52425 Jülich, Germany

**Keywords:** Evolution, Fungi, Genetic modification, Genomics, Lifestyle, Powdery mildew

## Abstract

Obligate biotrophic plant pathogens like the powdery mildew fungi commit to a closely dependent relationship with their plant hosts and have lost the ability to grow and reproduce independently. Thus, at present, these organisms are not amenable to *in vitro* cultivation, which is a prerequisite for effective genetic modification and functional molecular studies. Saprotrophic fungi of the family *Arachnopezizaceae* are the closest known extant relatives of the powdery mildew fungi and may hold great potential for studying genetic components of their obligate biotrophic lifestyle. Here, we established telomere-to-telomere genome assemblies for two representatives of this family, *Arachnopeziza aurata* and *A. aurelia*. Both species harbor haploid genomes that are composed of 16 chromosomes at a genome size of 43.1 and 46.3 million base-pairs, respectively, which, in contrast to most powdery mildew genomes that are transposon-enriched, show a repeat content below 5% and signs of repeat-induced point mutation (RIP). Both species could be grown in liquid culture and on solid standard media and were sensitive to common fungicides such as hygromycin and fenhexamid. We successfully expressed a red fluorescent protein and hygromycin resistance in *A. aurata* following polyethylene glycol-mediated protoplast transformation, demonstrating that *Arachnopeziza* species are amenable to genetic alterations that may include gene replacement, gene modification, and gene complementation. With this work, we established a potential model system that promises to sidestep the need for genetic modification of powdery mildew fungi by using *Arachnopeziza* species as a proxy to uncover the molecular functions of powdery mildew proteins.

## Introduction

Powdery mildew is one of the most important and widespread plant diseases, visible as a greyish-white tarnish or pustules on infected plant tissues such as leaves and stems (Glawe, 2008). The disease is caused by the ascomycete powdery mildew fungi (family of *Erysiphaceae*), which comprise 19 genera and around 900 species infecting more than 10,000 angiosperm plant species (Braun & Cook, 2012; Kiss *et al*., 2020; Kusch *et al*., 2024b). The powdery mildews are obligate biotrophic plant pathogens, meaning they are fully dependent on living host plant tissue for growth and reproduction (Kemen *et al*., 2015; Spanu & Panstruga, 2017). This lifestyle is associated with genomic hallmarks, including genome size expansion and gene losses in primary and secondary metabolism (Spanu *et al*., 2010; Frantzeskakis *et al*., 2019). The genome size of currently available near-chromosome assemblies of powdery mildew fungi ranges from 76-212 million base pairs (Mbp), which is more than two times above the average ascomycete genome size of around 37 Mbp (Mohanta & Bae, 2015; Kusch *et al*., 2024b). The only known exception is *Parauncinula polyspora*, a powdery mildew with a comparatively small (< 30 Mbp) genome that represents the most ancient extant lineae within the *Erysiphaceae* (Frantzeskakis *et al*., 2019; Vaghefi *et al*., 2022). In all other species with known genome assemblies, the size expansion can be attributed largely to transposable elements (TEs), which frequently cover 60-80% of the genomes in powdery mildew fungi (Spanu *et al*., 2010; Frantzeskakis *et al*., 2018; Müller *et al*., 2019; Kusch *et al*., 2024b). Meanwhile, genes encoding components required for the assimilation of nitrogen and sulfur, anaerobic fermentation, and thiamine biosynthesis are missing in the powdery mildew genomes, signifying their dependence on the host to acquire important nutrients and metabolites (Spanu *et al*., 2010; Frantzeskakis *et al*., 2019). As a consequence, it is presently not possible to cultivate or genetically modify powdery mildew fungi. Although alternative approaches such as host-induced gene silencing (Nowara *et al*., 2010), mutagenesis approaches coupled with selection (Barsoum *et al*., 2020; Bernasconi *et al*., 2024), and experimental evolution (Schwarzbach, 1979; Kusch *et al*., 2024a) have been used successfully for studying aspects of powdery mildew virulence and evolution, the lack of cultivation and genetic modification protocols severely limits the functional molecular and genetic analysis of the obligate biotrophic lifestyle.

Based on multi-gene and nuclear ribosomal DNA (nrDNA) phylogenetic analyses (Johnston *et al*., 2019; Vaghefi *et al*., 2022), the closest extant relatives of the *Erysiphaceae* known to date are members of the family *Arachnopezizaceae*. Five genera are currently assigned to the *Arachnopezizaceae*. The genera *Arachnopeziza*, *Arachnoscypha*, and *Eriopezia* have been included in the *Arachnopezizaceae* based on molecular phylogenetic analysis (Han *et al*., 2014), although *Arachnoscypha* was recently suggested to belong to the family *Hyaloscyphaceae* (Kosonen *et al*., 2020). Additionally, *Austropezia* and *Parachnopeziza* are assigned to the *Arachnopezizaceae* based on morphological features (Han *et al*., 2014; Baral, 2015). The genus *Arachnopeziza* was first established in 1870 with the type species *A. aurelia* (Fuckel, 1870), but *A. aurata* was later suggested to be the type species (Korf, 1951). Currently, 12 (Kosonen *et al*., 2020) to 15 (Kirk *et al*., 2008) species are recognized within the genus *Arachnopeziza*, but several undescribed species are likely to exist.

The *Arachnopezizaceae* are saprobic fungi from the phylum Ascomycota and comprise species living on dead plant matter such as wood and litter (Korf, 1951; Ekanayaka, 2019). These fungi have thick-walled hyaline hyphae. The asexual morph is hyphomycetous, i.e., asexual spores (conidia) are formed from hyphae. During sexual reproduction, the fungi develop septate ellipsoid to fusoid ascospores (Korf, 1951; Kirk *et al*., 2008; Ekanayaka, 2019). The fruiting bodies are formed on a subiculum, which is an interconnected web-like mycelium established by protruding hyphal elements and surrounding the apothecia (Kirk *et al*., 2008). The subiculum together with septate spores are shared phenotypic characteristics of the *Arachnopezizaceae*, distinguishing them from other closely related families (Korf, 1951; Han *et al*., 2014; Kosonen *et al*., 2020). Several *Arachnopeziza* species appear to be intimately associated with mosses such as *Sphagnum* and liverworts like *Ptilidium*, but the biological relevance of this relationship remains unclear (Stenroos *et al*., 2010; Kosonen *et al*., 2020).

Because the *Arachnopezizaceae* are phylogenetically the closest known extant relatives of the powdery mildew fungi (Johnston *et al*., 2019; Vaghefi *et al*., 2022) and exhibit a non-pathogenic saprobic lifestyle (Korf, 1951), we hypothesized that these fungi could constitute a suitable experimental system to establish molecular tools for genetic studies. We selected two species, *A. aurata* and *A. aurelia*, each represented by a strain deposited as a living culture at the CBS-KNAW fungal culture collection under accession numbers CBS127674 and CBS127675, respectively. We generated chromosome-level genome assemblies and gene annotation resources, worked out cultivation conditions, and established a protoplast-mediated transformation protocol, highlighting the potential of these fungal species to facilitate future molecular studies related to the obligate biotrophic lifestyle of the powdery mildew fungi.

## Results

### *Arachnopeziza* species grow under *in vitro* culture conditions

Effective and reproducible genetic modification protocols rely on *in vitro* cultivation methods (Lichius *et al*., 2020). We tested if *A. aurata* CBS127674 and *A. aurelia* CBS127675 can be cultivated on standard media used for cultivation of fungi, i.e., maltose extract agar (MEA), potato dextrose agar (PDA), and yeast peptone dextrose agar (YPDA). Both fungi formed visible mycelium on the three media but did not grow on lysogeny broth (LB), a standard growth medium for bacteria (**Figure 1A**). Fungal growth was detected at 23 °C and 28 °C (**Figure 1B**) but was restricted at 37 °C, and an increase in humidity by the addition of sterile water on PDA or MEA positively affected the colony size of both fungal species (**Figure 1C**). Both fungi also grew well in liquid medium (potato dextrose broth; PDB) at 28 °C within 7 days of cultivation (**Figure 1D**). In addition, we were able to recover growing mycelium of *A. aurata* but not *A. aurelia* after flash-freezing of liquid culture samples in glycerol at concentrations ranging from 10-30% after up to one year of storage at -80 °C (**Figure 1E**). As *Arachnopeziza* species are wood- or plant litter-decaying fungi (Korf, 1951; Ekanayaka, 2019), we also added sterilized toothpicks or pieces of twigs and leaves from a local maple tree to MEA. We found that *A. aurelia* specifically overgrew leaves but not wood, while *A. aurata* overgrew wood but not leaves after 3 months of cultivation (**Figure 1F**). Overall, we successfully cultivated two *Arachnopeziza* species *in vitro* using media commonly used for fungal propagation.

**Figure 1.**
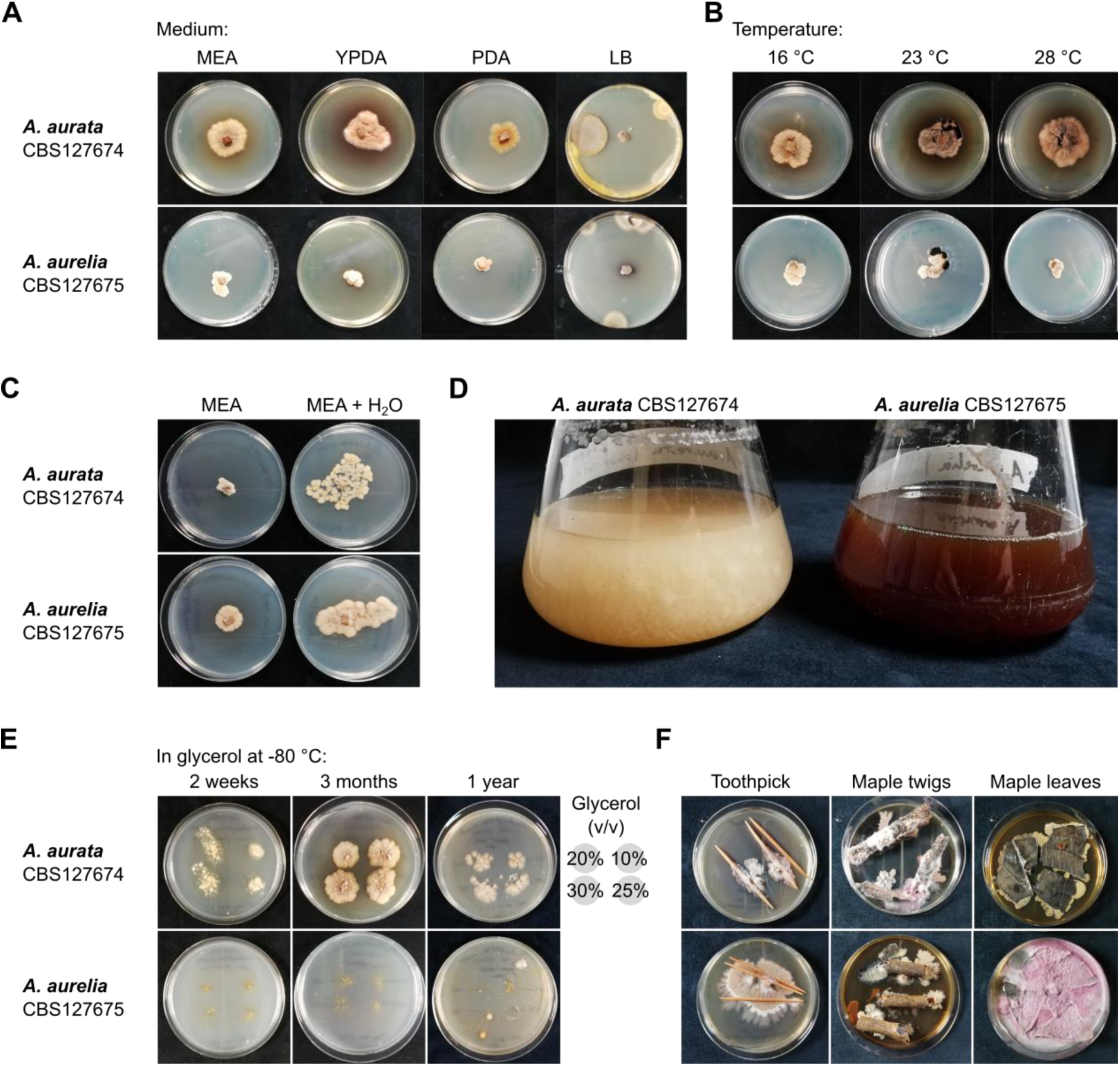
*Arachnopeziza aurata* and *A. aurelia* can be cultivated under standard *in vitro* growth conditions. The strains *Arachnopeziza aurata* CBS127674 and *A. aurelia* CBS127675 were cultivated on solid and liquid medium, inoculated using agar plugs containing mycelium. (**A**) *A. aurata* and *A. aurelia* were incubated on agar plates (from left to right, maltose extract agar (MEA), yeast peptone dextrose agar (YPDA), potato dextrose agar (PDA), and lysogeny broth (LB) at 23 °C. Photos were taken 24 days after inoculation. (**B**) The fungi were grown on MEA plates at 16 °C, 23 °C, or 28 °C. Photos were taken 24 days after inoculation. (**C**) *A. aurata* and *A. aurelia* were incubated on dry MEA plates (left) or by the addition of 200 µL of sterile H_2_O at 23 °C. Photographs were taken 14 days after inoculation. (**D**) The fungi were grown in potato dextrose broth (PDB) at 80 revolutions per minute (rpm) and 28 °C. Photos taken 12 days after inoculation. (**E**) *A. aurata* and *A. aurelia* were incubated in PDB, and glycerol was added at final concentrations of 10% (v/v), 20% (v/v), 25% (v/v), and 30% (v/v), arranged as indicated on each plate; cultures were then flash-frozen in liquid nitrogen and stored at - 80 °C. Cultures were recovered after 2 weeks, 3 months, and 1 year of storage by inoculating 50 µL of glycerol stock on MEA and incubation at 23 °C for 14 days. (**F**) MEA plates were covered with (from left to right) sterile toothpicks, sterile maple twigs, or sterile maple leaf segments and inoculated with *A. aurata* or *A. aurelia*. The plates were incubated at 23 °C, and photographs were taken after 3 months.

### *Arachnopeziza* species have compact genomes

To generate genomic resources for the fungal species *Arachnopeziza aurata* and *A. aurelia*, we obtained long sequence reads using MinION nanopore technology from *in vitro*-cultured fungi for genome assembly. We merged the assemblies generated with three long-read assemblers (Canu, Flye, and NextDenovo), yielding haploid genomes composed of 16 nuclear pseudo-chromosomes for both species (**Table 1**, **Figure 2A**). We then used teloclip (https://github.com/Adamtaranto/teloclip) to retrieve nanopore reads at both ends of the assembled sequences containing telomeric repeats (5’-TTAGGG-3’) and thus recovered 31 of the 32 telomeric ends in the genome assembly of *A. aurata* and 30 out of 32 telomeres in the case of *A. aurelia* (**Figure 2A**). Additionally, we determined putative centromeric regions of the 16 pseudo-chromosomes per species. The final assemblies contained 43,078,696 base pairs (bp) in the case of *A. aurata* and 46,346,353 bp for *A. aurelia* with N50 values of 2,695,373 bp and 2,924,875 bp, respectively (**Table 1**). To confirm the identity of both assemblies, we extracted the 5.8S, 18S, 28S nuclear ribosomal DNA (nrDNA) and the corresponding internal transcribed spacer (ITS) sequences and found that they were 99% identical to the respective nrDNA sequences deposited in GenBank (accessions MH864617.1, MH876055.1, MH864618.1, and MH876056.1; see **Supplementary Files 1** and **2** for alignments). We also used Benchmarking Universal Single-Copy Orthologs (BUSCO) to assess the completeness of the two genome assemblies and found 99.0% of the 1,706 core ascomycete genes to be present as single copies in the two assemblies (**Table 1**). By comparison, the available draft assembly of *A. araneosa* (Johnston *et al*., 2019) was generated using short-read sequencing from an environmental sample and, therefore, is highly fragmented, as e.g. indicated by a low N50 value of 210,669 bp (more than 10-times lower than for *A. aurata* and *A. aurelia*; **Table 1**). Thus, the assemblies generated in the current work comprise significantly improved resources for *Arachnopeziza* species. The genomes of the two species *A. aurata* and *A. aurelia* exhibited overall high levels of collinearity, although structural rearrangements could be observed for some chromosomes (**Figure 2B**). For instance, *A. aurelia* chromosome 6 was collinear in part with *A. aurata* chromosomes 1 and 11.

**Figure 2.**
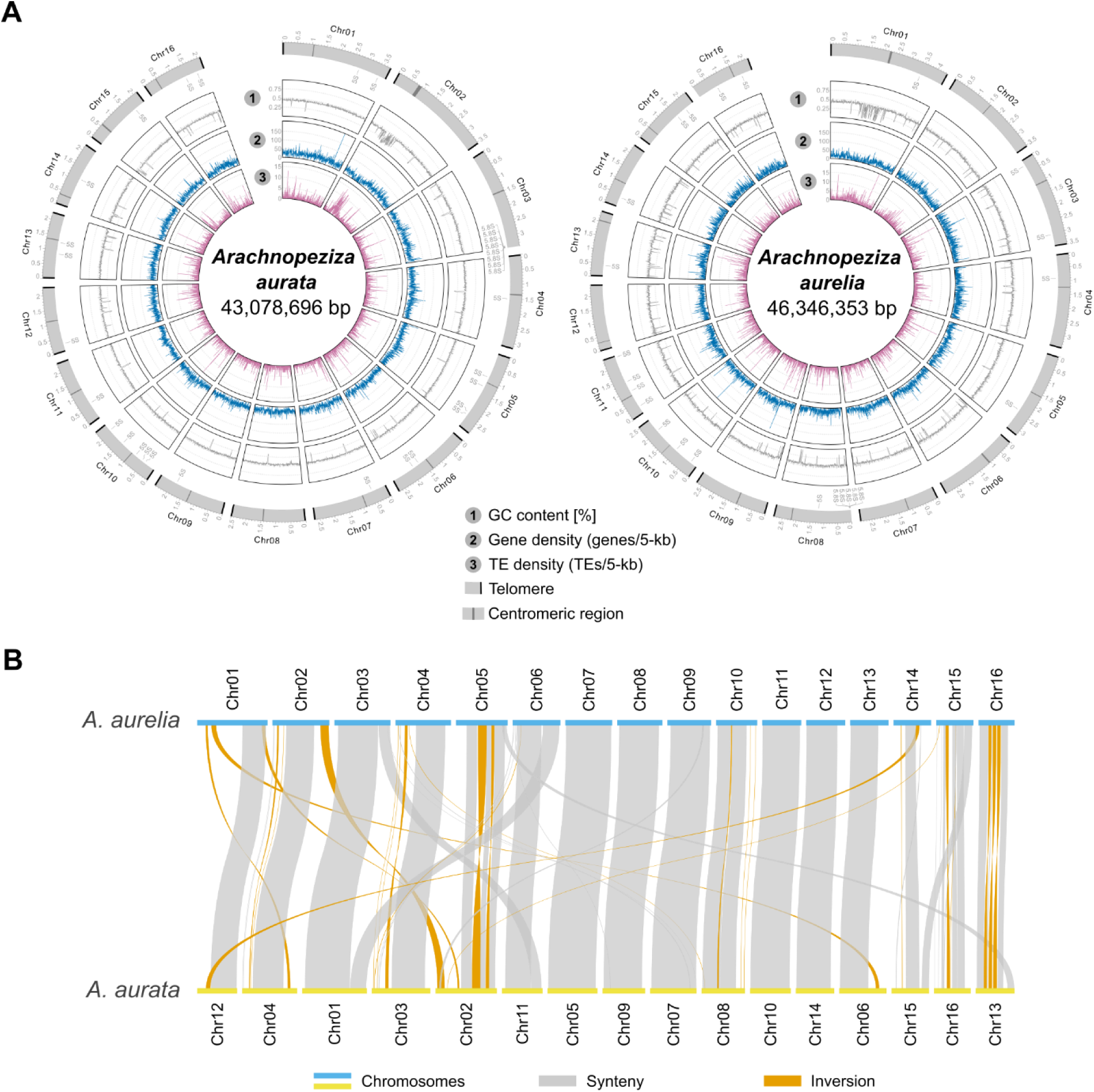
The chromosome-scale genome assemblies for *Arachnopeziza aurata* and *A. aurelia* exhibit a high level of synteny. We generated genome assemblies for *A. aurata* CBS127674 and *A. aurelia* CBS127675 using MinION nanopore technology. (**A**) The circos plots display the genomic maps for *A. aurata* (left panel) and *A. aurelia* (right panel) and were generated with the R package circlize v0.4.10 (Gu *et al*., 2014). The outer tracks indicate the chromosomal map and the scale the position in million base pairs (Mbp); black rectangles denote telomeres and dark grey rectangles putative centromeres. We searched for potential candidate centromeric regions by identifying sliding windows covering between 30,000-150,000 bp where the gene content was zero or near zero and the TE content was at least 5 TEs per window. The tracks show GC content (1), gene density expressed as the number of genes per 5-kb window (2), and TE density calculated as TEs per 5-kb window (3) as line graphs. (**B**) The synteny plot shows the collinearity between the genome assemblies of *A. aurata* and *A. aurelia* and was generated with the R package RIdeogram v0.2.2 (Hao *et al*., 2020). The blue and yellow lines display the respective chromosomes, grey bars show blocks of synteny, and orange bars denote inversions.

**Table 1.** Genome assembly statistics for *A. aurata* and *A. aurelia* compared to *A. araneosa*.

| Feature | *A. araneosa*<sup>a</sup> | *A. aurata* | *A. aurelia* |
| --- | --- | --- | --- |
| Genome assembly [bp] | 40,453,245 | 43,078,696 | 46,346,353 |
| N count | 3,840 | 0 | 0 |
| Estimated size (k-mer) [bp] | - | 43,965,082 | 45,594,330 |
| Scaffolds/contigs | 1,498 | 16 | 16 |
| Largest sequence [bp] | 1,164,973 | 3,758,135 | 4,390,527 |
| Average length [bp] | 27,004.8 | 2,692,418.5 | 2,896,647.1 |
| GC content [%] | 45.63 | 44.49 | 43.26 |
| N50 | 210,669 | 2,695,373 | 2,924,875 |
| N90 | 46,250 | 2,227,591 | 2,344,259 |
| CRAQ<sup>b</sup> |  |  |  |
| Sequence covered [%] | - | 99.99 | 99.99 |
| R-AQI | - | 92.84 | 93.32 |
| S-AQI | - | 100.00 | 100.00 |
| BUSCO<sup>c</sup> |  |  |  |
| Complete single-copy | 1,690 (99.1%) | 1,689 (99.0%) | 1,688 (99.0%) |
| Duplicated | 2 (0.1%) | 3 (0.2%) | 2 (0.1%) |
| Fragmented | 12 (0.7%) | 13 (0.8%) | 15 (0.9%) |
| Missing | 2 (0.1%) | 1 (0.0%) | 1 (0.0%) |
| Interspersed repeats [%]<sup>d</sup> | 0.83 | 2.63 | 3.93 |
| SINE retroelements [%] | 0.00 | 0.01 | 0.00 |
| LINE retroelements [%] | 0.00 | 0.00 | 0.00 |
| LTR elements [%] | 0.72 | 2.36 | 2.50 |
| DNA transposons [%] | 0.11 | 0.20 | 0.10 |
| Other/Unclassified [%] | 0.01 | 0.07 | 1.33 |

<sup>a</sup> Strain D781; retrieved from NCBI genomes database at accession ASM398885v1 (Johnston *et al.*, 2019)
<sup>b</sup> Determined using CRAQ v1.0.9 (Li *et al.*, 2023); R-AQI, regional assembly quality index; S-AQI, overall assembly quality index. AQI values > 90 are considered reference quality.
<sup>c</sup> Ran with 1,706 core ascomycete genes from the database ascomycota\_odb10 using compleasm version 0.2.6 (Simão *et al.*, 2015; Huang & Li, 2023)
<sup>d</sup> Database was calculated with an EDTA2-TETrimmer pipeline (Qian *et al.*, 2024)

In addition, we assembled the mitochondrial genomes of both species. The mitochondrial genomes had a size of 28,111 bp (*A. aurata*) and 28,095 bp (*A. aurelia*), respectively. By comparison, currently available fungal mitogenomes range from 12 kb in *Rozella allomycis* (James *et al*., 2013) to 531 kb in *Morchella crassipes* (Liu *et al*., 2020). All components of the mitochondrial respiratory complexes I, III, and IV, both small and large ribosomal RNAs *rns* and *rnl*, the gene coding for ribosomal protein S3 (*rps3*), and the ribonucleoprotein RNA *rnpB*, which are typically found in fungal mitochondrial genomes (Seif *et al*., 2005; Aguileta *et al*., 2014), were present. The sole exception was the ATP synthase component *atp8*, which we did not detect in either of the two mitochondrial genomes (**Supplementary Figure 1**). Loss of *atp8* (along with *atp9*) in the mitochondrial genomes due to migration to the nuclear genome has been observed for other fungi, such as *Stemphylium lycopersici* (Franco *et al*., 2017). Similarly, we found an orthologue of fungal *atp8* in each of the nuclear genomes of *A. aurata* and *A. aurelia*, encoded by *AAURAT_014873-RA* on Chr15 and *AAUREL_012968-RA* on Chr14, respectively. Thus, *atp8* migrated to the nuclear genome in the two *Arachnopeziza* species. By contrast, *atp8* is present in the mitochondrial genomes of powdery mildew fungi such as *Erysiphe necator* (Zaccaron *et al*., 2021), *B. graminis* f. sp. *tritici*, *E. pisi*, and *Golovinomyces cichoracearum* (Zaccaron & Stergiopoulos, 2021).

Next, we used EDTA2 and TEtrimmer (Ou *et al*., 2019; Qian *et al*., 2024) to annotate transposable elements (TEs) and repeats in the genomes of *A. aurata* and *A. aurelia*. The species exhibited a repeat content of 2.6% and 3.9% (**Table 1**), respectively, which is well below the values of most powdery mildew fungi, which typically have repeat contents of 40-80% (Kusch *et al*., 2024b). The transposon space was dominated by long terminal repeat (LTR) retroelements, while long interspersed nuclear elements (LINEs) were absent in both genomes (**Table 1**). TEs were mostly located in gene-sparse regions with low GC content (**Figure 2A**), consistent with a general AT-richness of TE-derived open reading frames (Jia & Xue, 2009) and observations that TEs often cluster in gene-poor genomic regions in, for instance, the plant pathogen *Zymoseptoria tritici* (Oggenfuss & Croll, 2023). In contrast to powdery mildew fungi, whose genomes often show signatures of recent or ongoing TE bursts and no signs of the fungal TE defense mechanism repeat-induced point mutation (RIP) (Frantzeskakis *et al*., 2018, 2019; Kusch *et al*., 2024b), the TEs in the *Arachnopeziza* genomes had a peak sequence divergence of 2-10%, suggestive of evolutionary older transposition events, and exhibited weak but detectable bipartite GC content distribution, indicative of the RIP defense mechanism (**Figure 3A** and **3B**). We performed RIP index analysis using dinucleotide ratios calculated by RIPCAL (Hane & Oliver, 2008) and compared the *Arachnopeziza* genome assemblies with that of the barley powdery mildew pathogen *Blumeria hordei* DH14, which does not exhibit RIP signatures (Frantzeskakis *et al*., 2018), and *Rhynchosporium commune*, an example for a genome exhibiting enrichment of AT-rich sequences in repetitive regions, indicative of RIP activity (Frantzeskakis *et al*., 2019) (**Figure 3C**; **Supplementary Figure 2**). In the prototypical case of the model fungus *Neurospora crassa*, sequences with a TpA/ApT index above 0.89 and a (CpA + TpG)/(ApC + GpT) index below 1.03 suggest AT enrichment due to RIP (Margolin *et al*., 1998). We found the TpA/ApT indices of *Ty1*/*Copia* and *Ty3*/*mdg4* LTR retroelements to be above 1.9 in the case of *A. aurata* and above 1.3 in the case of *A. aurelia*, while the index was above 1.3 for DNA transposons in both species (**Figure 3C**). By comparison, the TpA/ApT ratio was around 0.6 for coding sequences (CDSs) and around 0.8 genome-wide. On the other hand, the (CpA + TpG)/(ApC + GpT) indices were below 0.5 for both types of LTR retrotransposons in both species, while DNA elements had an index of 0.63 for *A. aurelia* and 1.09 for *A. aurata*; the indices for CDSs were 1.35 and genome-wide 1.25 (**Figure 3C**). The difference between CDSs and the analyzed TEs was even more pronounced for the (CpA + TpG)/TpA index. The pattern of markedly increased TpA/ApT in combination with decreased (CpA + TpG)/(ApC + GpT) and (CpA + TpG)/TpA indices in DNA transposons and retroelements was similar in *R. commune*, while these indices exhibited no discernible difference for all analyzed features in *B. hordei*, reflecting the observed presence and absence, respectively, of RIP in these organisms (Frantzeskakis *et al*., 2018, 2019). Furthermore, we detected one orthologue each of the key components of RIP in fungi, i.e., *Masc1*, *Masc2*, *Rid-1*, and *Dim-2* (GenBank accession numbers AAC49849.1, AAC03766.1, XP_011392925.1, and XP_959891.1, respectively) (Gladyshev, 2017), in *A. aurata* (E values 5e-63, 1e-70, 8e-89, and 0.0, respectively), and one orthologue of *Rid-1* (E value 2e-78) and *Dim-2* (E value 0.0) each in *A. aurelia* (**Supplementary File 3** and **4**), while these genes are known to be absent in powdery mildew fungi (Spanu *et al*., 2010; Frantzeskakis *et al*., 2019; Kusch *et al*., 2024b). Overall, the genome assemblies of both *A. aurata* and *A. aurelia* showed an enrichment of AT dinucleotide signatures in LTR retroelements and, to a different degree, in DNA transposons, in combination with the presence of key RIP components, indicative of a functional RIP defense mechanism in these fungi.

**Figure 3.**
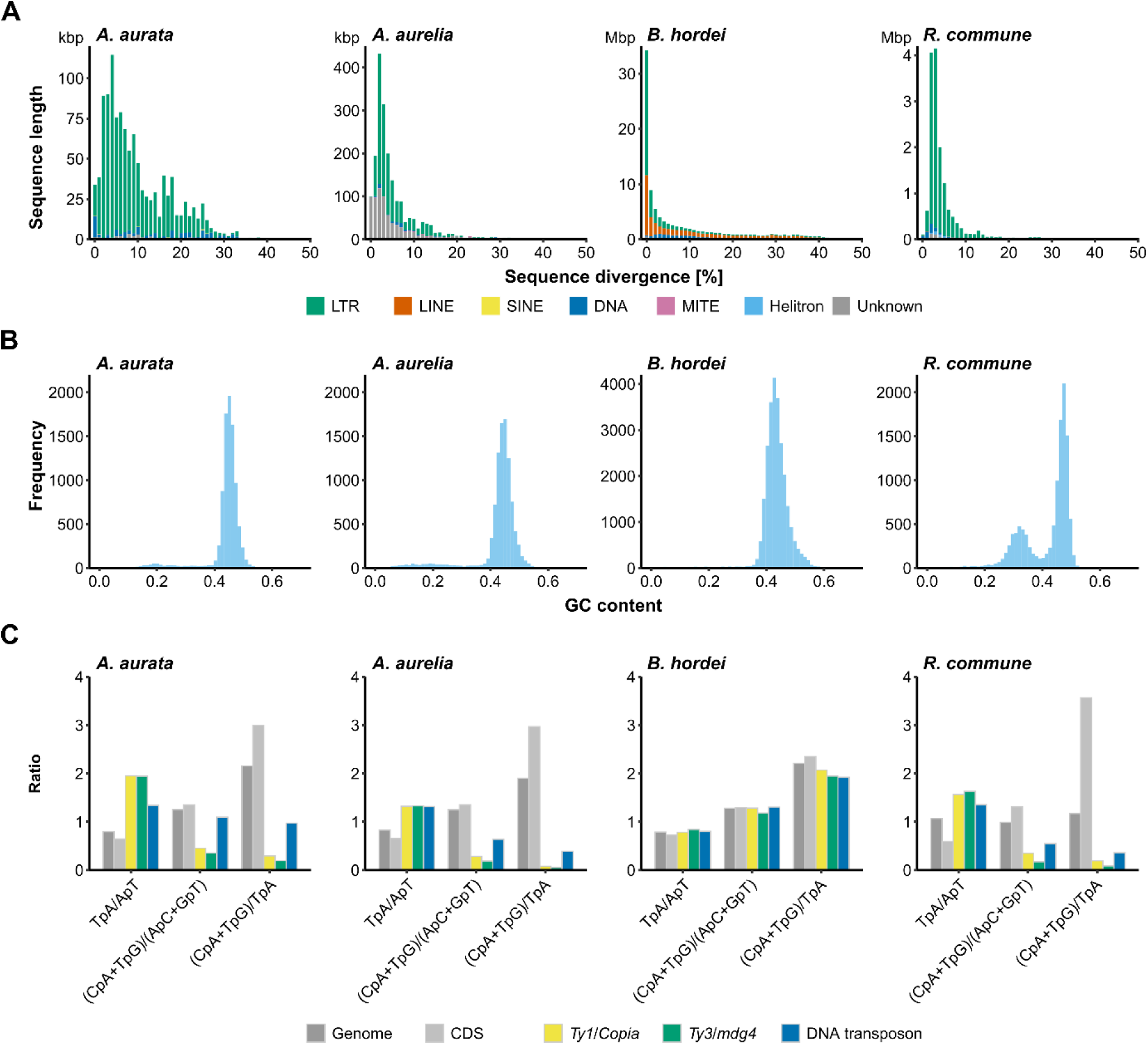
The genomes of *Arachnopeziza aurata* and *A. aurelia* show signs of TE degeneration. We compared the repetitive content of the genome assemblies of *A. aurata* CBS127674 and *A. aurelia* CBS127675 with the assemblies of the barley powdery mildew *Blumeria hordei* DH14 (Frantzeskakis *et al*., 2018) and the barley leaf scald disease pathogen *Rhynchosporium commune* UK7 (Penselin *et al*., 2016). Panels are displayed in this order from left to right. (**A**) The stacked bar graphs show the sequence divergence of TE subclasses in percent (x-axis) and the cumulative length of sequence at the respective level of divergence in base pairs (kbp or Mbp, as indicated; y-axis). The bars are colored according to TE class: long terminal repeat (LTR), green; long interspersed nuclear element (LINE), orange; short interspersed nuclear element (SINE), yellow; DNA transposons, blue; miniature inverted-repeat transposable element (MITE), purple; DNA Helitron, light blue; unknown or unclassified, grey. (**B**) The histograms display the distribution of GC content (ratio, x-axis) determined in genome-wide 5-kb windows; the y-axis shows the frequency of the respective content bin. (**C**) The bar graphs show the ratios of repetitive sequence dinucleotide analysis genome-wide (dark grey), for coding sequences (CDS; light grey), *Ty1*/*Copia* LTR elements (yellow), *Ty3*/*mdg4* LTR elements (orange), and DNA transposons (blue). The respective RIP index represented by dinucleotide ratios is indicated on the x-axis, and the y-axis displays the respective ratio.

### *Arachnopeziza* species exhibit the genomic signatures of a saprobic fungus with plant cell wall-degrading ability

We used global RNA sequencing data obtained from mycelium grown in liquid culture for both species (**Figure 1D**) to discover and annotate coding genes using BRAKER3 (Gabriel *et al*., 2021, 2023; Bruna *et al*., 2023). Overall, we predicted 15,452 genes in *A. aurata* and 13,437 genes in *A. aurelia* (**Table 2**), which is higher than the average gene number of ∼11,000 in ascomycetes (Mohanta & Bae, 2015).

**Table 2.** Gene annotation of *A. aurata* and *A. aurelia*.

| Feature | *A. aurata* | *A. aurelia* |
| --- | --- | --- |
| Number of coding genes <sup>a</sup> | 15,452 | 13,437 |
| Putatively secreted proteins <sup>b</sup> | 1,233 | 1,109 |
| Putative effectors <sup>c</sup> | 536 | 433 |
| Predicted CAZymes <sup>d</sup> | 618 | 503 |
| Contains protein family (PFAM) domain | 12,360 | 10,505 |
| Contains InterPro Superfamily domain | 8,049 | 6,831 |
| Secondary metabolism component <sup>e</sup> | 69 | 73 |
| Non-ribosomal peptide synthetases (NRPS) | 14 | 25 |
| Polyketide synthases (PKS) | 31 | 22 |
| Fungal ribosomally synthesized and post-translationally modified peptide (RiPP)-like | 18 | 21 |
| Terpene biosynthesis | 6 | 4 |
| Indole biosynthesis | 0 | 1 |

<sup>a</sup> Genes predicted using BRAKER3 (Gabriel *et al.*, 2021, 2023; Bruna *et al.*, 2023)
<sup>b</sup> Secretion signals were predicted using SignalP5.0 (Almagro Armenteros *et al.*, 2019)
<sup>c</sup> Predicted using EffectorP3.0 (Sperschneider & Dodds, 2021)
<sup>d</sup> CAZymes were predicted using dbCAN3 (Zheng *et al.*, 2023)
<sup>e</sup> Predicted using antiSMASH v7.1.0 (Medema *et al.*, 2011; Blin *et al.*, 2023)

We used Kyoto Encyclopedia of Genes and Genomes (KEGG) Mapper (https://www.kegg.jp/) to reconstruct the primary metabolism pathways in *A. aurata* and *A. aurelia* (**Figure 4A**, **Supplementary Table 1**, **2**, and **3**). We recovered most metabolic pathways related to carbohydrate, energy, lipid, amino acid, and cofactor metabolism in *A. aurata*. In the case of *A. aurelia*, several components of the amino acid metabolism were not found, specifically of the branched-chain amino acid metabolism pathways related to valine, isoleucine, and leucine biosynthesis. Nonetheless, both fungi display a comparable primary metabolism profile to the fungal plant pathogen *Botrytis cinerea* (**Figure 4A**) and do not show the losses in sulfur assimilation and thiamine biosynthesis reported for the obligate biotrophic powdery mildew fungus *B. hordei* (Spanu *et al*., 2010; Frantzeskakis *et al*., 2018) and other powdery mildew species (Frantzeskakis *et al*., 2019).

**Figure 4.**
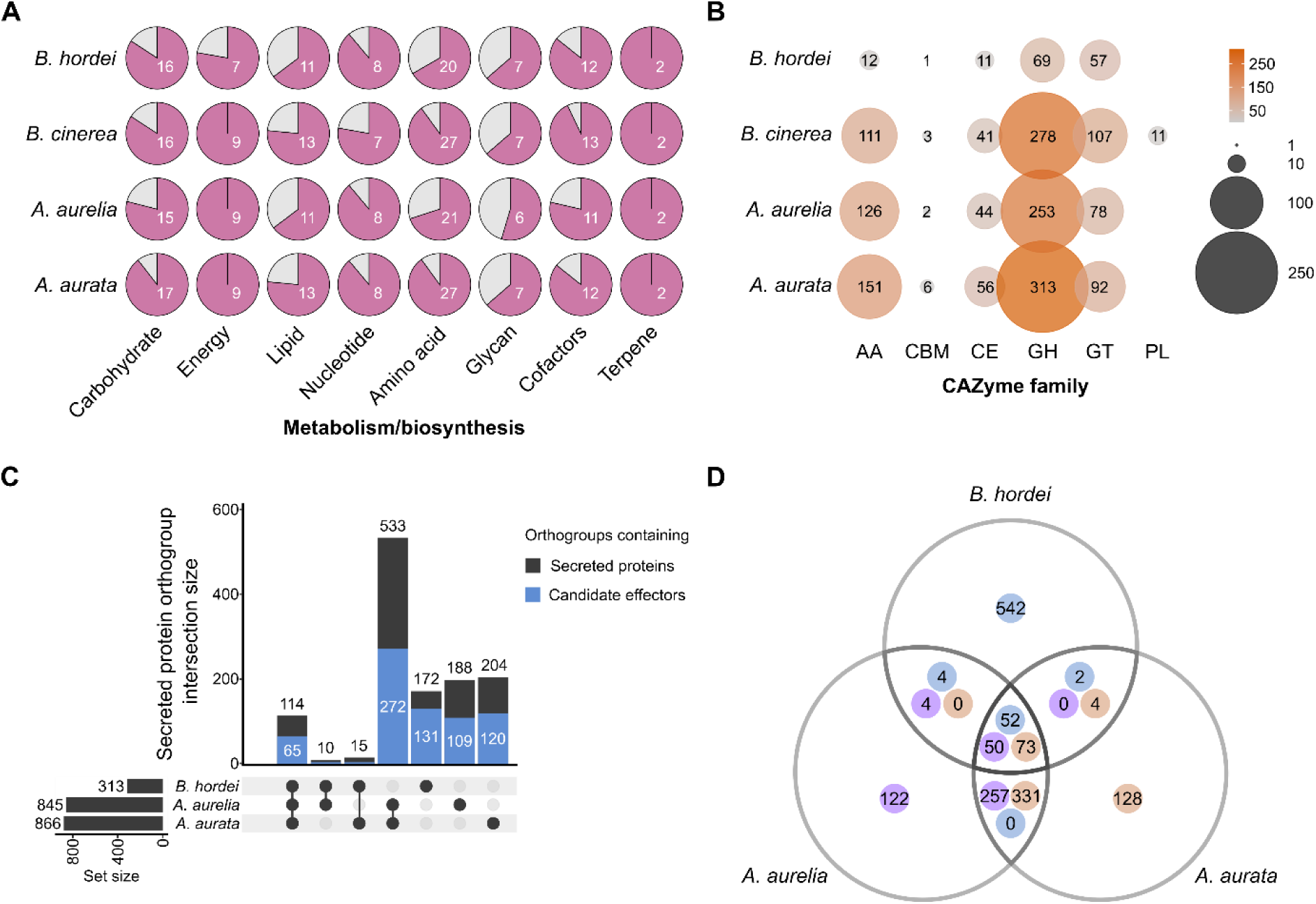
***Arachnopeziza aurata* and *A. aurelia* harbor genes enabling a saprobic lifestyle and plant cell wall degradation and possess candidate effectors that may facilitate host interactions.** (A) Primary metabolism modules were annotated using GhostKOALA to identify components and KEGG Mapper to reconstruct pathways via https://www.kegg.jp/ (accessed 04/2025) in *A. aurata*, *A. aurelia*, *B. cinerea* B05.10 (Van Kan *et al*., 2017), and *B. hordei* DH14 (Frantzeskakis *et al*., 2018). The pie charts indicate the number of complete pathways in the respective categories (x-axis), i.e., carbohydrate metabolism (19 pathways in fungi), energy metabolism (9 pathways), lipid metabolism (17 pathways), nucleotide metabolism (9 pathways), amino acid metabolism (30 pathways), glycan metabolism (11 pathways), cofactor metabolism (14 pathways), and terpene biosynthesis (2 pathways). (B) We used dbCAN3 (Zheng *et al*., 2023) to annotate putative carbohydrate-active enzymes in *A. aurata*, *A. aurelia*, *B. cinerea* B05.10 (Van Kan *et al*., 2017), and *B. hordei* DH14 (Frantzeskakis *et al*., 2018). The dot plot indicates the number of genes annotated to encode accessory activity (AA) enzymes, carbohydrate-binding motif (CBM)-containing proteins, carbohydrate esterases (CE), glycosyl hydrolases (GH), glycosyl transferases (GT), and polysaccharide lyases (PL). The size of the dots and their shade of orange are proportional to the number of predicted genes in each category; the number is also written on each dot. (**C**) The UpSet plot shows the overlap of orthologous groups of putative secreted proteins from *B. hordei*, *A. aurata*, and *A. aurelia* (**Table 2**). Signal peptides for secretion were predicted using SignalP5.0 (Almagro Armenteros *et al*., 2019), and orthologous groups (orthogroups) were assigned with OrthoFinder v2.5.5 (Emms & Kelly, 2019). The blue portion of the bars indicates orthogroups containing at least one candidate effector protein as determined by EffectorP3.0 (Sperschneider & Dodds, 2021). The sets corresponding to each bar are indicated below the bar chart by black dots. (**D**) The Venn diagram illustrates the orthologous overlap of candidate effector proteins from *B. hordei* (blue dot), *A. aurata* (orange dot), and *A. aurelia* (purple dot), as determined by EffectorP3.0.

Further, 618 and 503 genes were predicted to encode carbohydrate-active enzymes (CAZymes) according to a dbCAN3 search (Zheng *et al*., 2023) (**Table 2**, **Supplementary Table 1** and **2**). Similar to the necrotroph *B. cinerea*, which is capable of digesting plant-derived carbohydrates including those present in cell walls (Kubicek *et al*., 2014), both *A. aurata* and *A. aurelia* possess large numbers of putative glycosyl hydrolases, glycosyl transferases, carbohydrate esterases, and auxiliary enzymes (**Figure 4B**). This contrasts the powdery mildew pathogen *B. hordei*, which exhibits a reduced CAZyme complement, particularly of glycosyl hydrolases and auxiliary enzymes (Frantzeskakis *et al*., 2018). Different from *B. cinerea*, the two *Arachnopeziza* species do not seem to harbor polysaccharide lyases, which cleave uronic acid-containing polysaccharide chains and include fungal pectin lyases (Lombard *et al*., 2010). The pectin lyase family appears to be expanded in plant pathogens but is often absent in saprobic fungi (Karlsson *et al*., 2015), consistent with the non-pathogenic lifestyle of the *Arachnopeziza* species.

According to analysis by SignalP5.0 (Almagro Armenteros *et al*., 2019), we found that 1,233 *A. aurata* and 1,109 *A. aurelia* genes are predicted to encode secreted proteins, of which 536 and 433 are putative effector proteins according to EffectorP3.0 (Sperschneider & Dodds, 2021) (**Table 2**). Of these, 266 (*A. aurata*) and 231 (*A. aurelia*) have unknown functions (no protein families (PFAM) domain detected; **Supplementary Table 1** and **2**). The most common functional predictions for the remaining effector candidates are related to carbohydrate-binding or modification, peptidase activity, and presence of a common fungal extracellular membrane (CFEM) domain. We compared the *Arachnopeziza* candidate effectors with annotated candidate secreted effector proteins (CSEPs) of *B. hordei* DH14 (Frantzeskakis *et al*., 2018) and added putative secreted proteins predicted to be effectors according to an EffectorP3.0 search (Sperschneider & Dodds, 2021), totaling 600 *B. hordei* candidate effectors. We found that 139 orthologous groups of proteins (orthogroups) are shared between *B. hordei*, *A. aurata*, and *A. aurelia* candidate secreted proteins, while 533 orthogroups are exclusively shared between the two *Arachnopeziza* species (**Figure 4C**; **Supplementary Table 4**). Of the 139 shared orthogroups, 65 contained at least one candidate effector. Among these, 52 *B. hordei* candidate effectors were orthologous with 73 *A. aurata* and 50 *A. aurelia* candidate effectors (**Figure 4D**). On the other hand, 257 *A. aurelia* effectors were orthologous with 331 *A. aurata* candidate effectors, while these were absent in *B. hordei*. Likewise, most *B. hordei* candidate effectors (542) have no orthologue in either *Arachnopeziza* species, including the absence of seven powdery mildew core effectors (Sabelleck *et al*., 2025). The family of ribonuclease-like effectors expressed in haustoria (RALPHs) is ubiquitously present in powdery mildew fungi (Frantzeskakis *et al*., 2019) and particularly abundant in cereal powdery mildew fungi (Pedersen *et al*., 2012; Spanu, 2017). We searched the putative secretome of *A. aurata* and *A. aurelia* for the presence of domains of the ribonuclease superfamily (IPR016191) and found AAUREL_003350-RA as the only case of an effector candidate with this domain. Unlike RALPH effectors of powdery mildew fungi, AAUREL_003350-RA displays sequence conservation of the residues required for ribonuclease activity in RNase T1, suggesting that it is a catalytically active ribonuclease (**Supplementary Figure 3**; see **Supplementary File 5** for alignment). Additional searches for PFAM domains (PF00445, PF06479) and InterPro domains (IPR036430) from the ribonuclease T2-like superfamily identified three *A. aurata* and three additional *A. aurelia* secreted proteins that were assigned at least one of these domain accessions (**Supplementary Table 5**). However, sequence similarity analysis indicated that six of these ribonuclease-like proteins are more similar to fungal ribonuclease T1/T2/Trv proteins than RNase-like effectors of powdery mildew fungi (**Supplementary Figure 4**; see **Supplementary File 6** for alignment). One (AAURAT_002659) is an orthologue of the endoplasmic reticulum membrane-localized serine/threonine kinase inositol-requiring protein 1 (IRE1), which contains a ribonuclease domain and is involved in the unfolded protein (UPR) stress response (Credle *et al*., 2005). Overall, this analysis suggests that *Arachnopeziza* fungi do not harbor RNase-like effectors.

Fungi biosynthesize a large range of secondary metabolites to communicate and compete with other organisms, for protection from abiotic stress, and to regulate their development (Keller, 2019; Nguyen *et al*., 2023). We used antiSMASH v7.1.0 (Medema *et al*., 2011; Blin *et al*., 2023) to discover components of secondary metabolism in *A. aurata* and *A. aurelia*. Both genomes encode around 70 putative components of secondary metabolite biosynthesis, including non-ribosomal peptide synthetases (NRPS), polyketide synthase (PKS), and fungal ribosomally synthesized and post-translationally modified peptide-like (fungal-RiPP-like) (**Table 2**, **Supplementary Table 1** and **2**). Several of the genes encoding these components are located in close proximity and may constitute clusters involved in secondary metabolite biosynthesis (**Supplementary Figure 5**), a common feature found in fungi (Robey *et al*., 2021; Nguyen *et al*., 2023). By comparison, we found only eight candidate secondary metabolite biosynthetic genes in *B. hordei*, in line with observed gene losses in this fungus in the context of its obligate biotrophic lifestyle (Spanu *et al*., 2010; Frantzeskakis *et al*., 2018).

Taken together, we observed that the two *Arachnopeziza* species possess the genetic components required for a saprobic lifestyle, including the ability to generate secondary metabolites, possibly to govern microbial competition, communication, and fungal development. The large complement of CAZymes similar to that of the necrotrophic plant pathogen *B. cinerea*, which devours plant biomass, including cell walls, points to the ability to degrade plant cell wall components such as cellulose and lignin. Unexpectedly, we identified more than 400 effector candidates in the two fungi, suggesting that *Arachnopeziza* species are capable of interacting with unknown plant hosts and/or microbes.

### *A. aurata* and *A. aurelia* are sensitive to common fungicides and antibiotics

Generating genetically modified microbes requires efficient selection markers to isolate successfully modified individuals. Antibiotic and fungicide resistance are frequently used features that serve as selection markers in microbes. We used a panel of antibiotics and fungicides commonly deployed for the selection of transformants (**Table 3**) and cultivated *A. aurata* and *A. aurelia* on PDA containing these antimicrobial compounds at varying concentrations. The fungicides hygromycin B and fenhexamid and the antibiotic geneticin G418 exhibited growth-inhibiting activity starting at concentrations of 5-10 μg mL^-1^ and fully suppressed the growth of both fungi at 10-50 μg mL^-1^ (**Figure 5**). In addition, streptomycin showed some inhibitory capacity against *A. aurata* at 400 μg mL^-1^ but not against *A. aurelia* (**Supplementary Figure 6**). Further, kanamycin had inhibiting activity at 400 μg mL^-1^, while none of the other antibiotics affected the growth of the two *Arachnopeziza* species at the tested concentrations (**Figure 5**; **Supplementary Figure 6**). Hence, hygromycin B, fenhexamid, and geneticin qualify as effective selection markers for *A. aurata* and *A. aurelia*.

**Figure 5.**
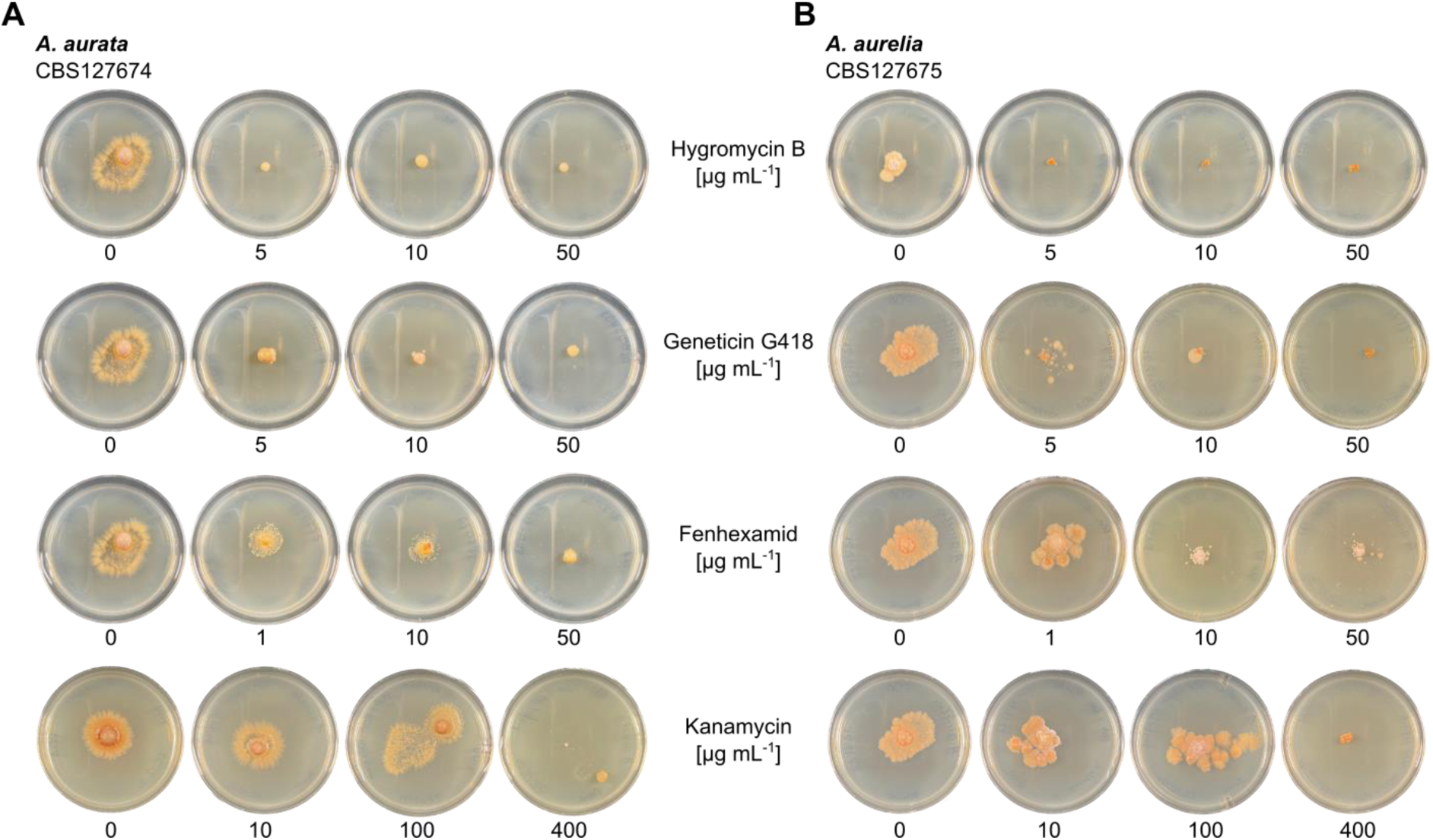
*Arachnopeziza aurata* and *A. aurelia* are inhibited by common fungicides. The strains (**A**) *Arachnopeziza aurata* CBS127674 and (**B**) *A. aurelia* CBS127675 were cultivated on potato dextrose agar (PDA) containing (top to bottom) hygromycin B, geneticin G418, fenhexamid, or kanamycin at the indicated concentrations. The plates were incubated at 23 °C; photographs were taken 16 days after inoculation. Additional fungicides and antibiotics, and the full set of tested concentrations for kanamycin and fenhexamid, are shown in **Supplementary Figure 6**. Note that the control pictures for the 0 μg mL^-1^ negative control are identical if the respective fungicides/antibiotics were tested in the same experiment.

### Protoplast-mediated transformation confers hygromycin resistance and fluorescent protein expression in *A. aurata*

Next, we aimed to determine if a common fungal transformation method can achieve genetic modification of *Arachnopeziza* species. We used a modified polyethylene glycol (PEG)-mediated protoplast transformation protocol established previously for *Magnaporthe oryzae* (Leisen *et al*., 2020; Wegner *et al*., 2022). After successful isolation of fungal protoplasts from liquid culture of *A. aurata* (**Figure 6A**), we used previously established vectors (pTelFen carrying the fenhexamid resistance allele of *Fusarium fujikuroi*, *FfERG27* (Cohrs *et al*., 2017; Wegner *et al*., 2022), and pTK144 conferring hygromycin resistance, and both harboring the monomeric red fluorescent protein (mRFP)-coding gene for protoplast transformation. We obtained one fenhexamid and one hygromycin B-resistant *A. aurata* line and confirmed the presence of the *FfERG27* (fenhexamid resistance allele), *HPH* (hygromycin resistance gene), and *mRFP* genes via genotyping by polymerase chain reaction (PCR) (**Figure 6B** and **6C**). We further confirmed the presence of the mRFP protein in the hyphae of some of the transformed lines by the detection of characteristic fluorescent signals via confocal laser scanning microscopy (**Figure 6D**). These experiments demonstrate that genes can be heterologously expressed in *A. aurata* using protoplast transformation.

**Figure 6.**
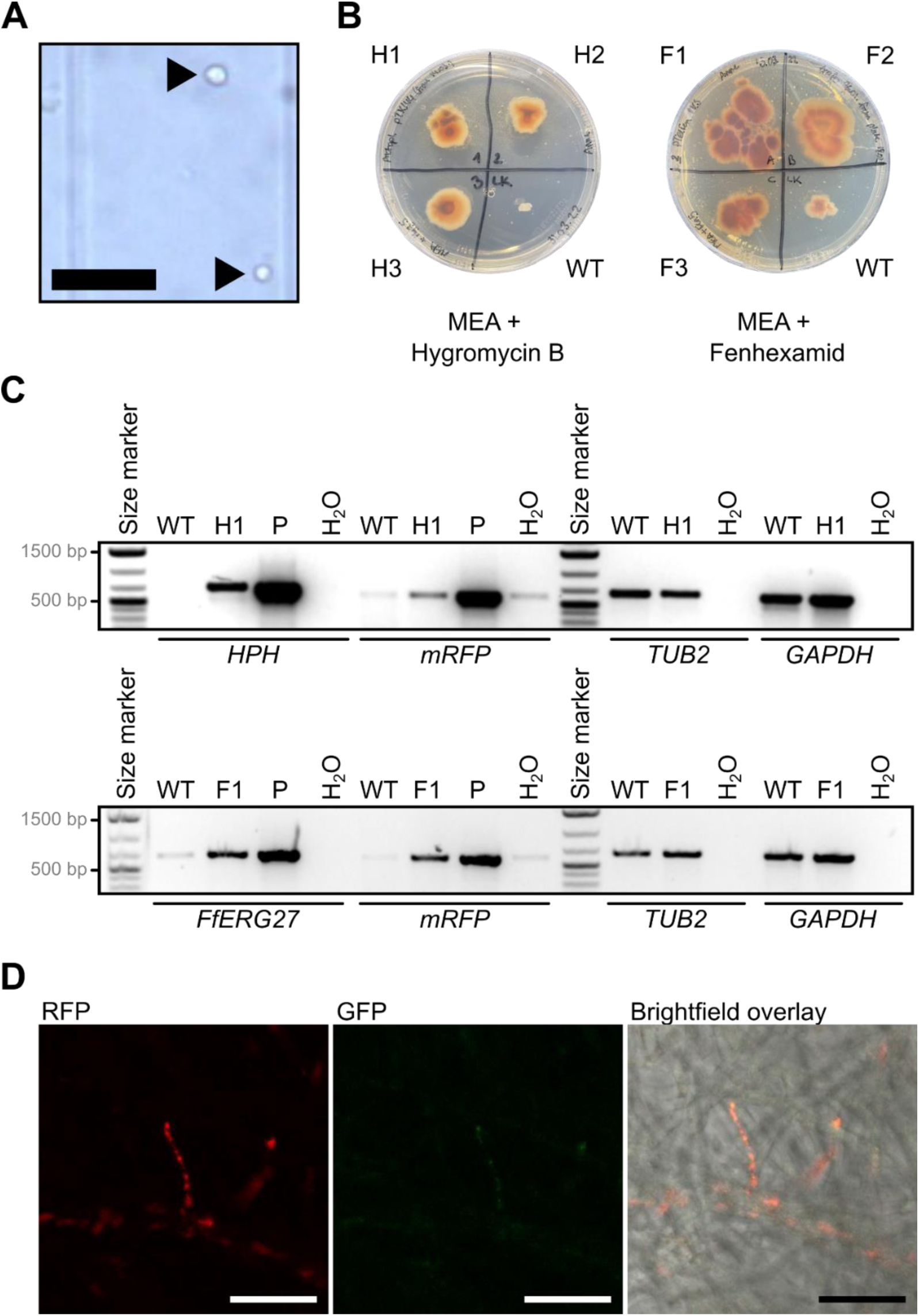
Transgenes conferring fungicide resistance and fluorescence can be expressed in *Arachnopeziza aurata*. *A. aurata* CBS127674 was genetically modified using a modified PEG-mediated protoplast transformation protocol (Leisen *et al*., 2020; Wegner *et al*., 2022). (**A**) Brightfield micrograph of *A. aurata* protoplasts (arrows); scale bar: 50 µm. (**B**) Strains were incubated on MEA with 5 µg mL^-1^ hygromycin B (left) or fenhexamid (right) at 23 °C. Transgenic strains are denoted H1-H3 (pTK144; hygromycin resistance) and F1-F3 (pTelFen, fenhexamid resistance). Photographs taken 16 days after incubation. (**C**) Genotyping PCRs for transgenic strains H1 and F1. The transgenes were: *HPH*, hygromycin resistance gene; *mRFP*, red fluorescent protein gene; *FfERG27*, fenhexamid resistance allele of *Fusarium fujikuroi*. Two endogenous *A. aurata* genes were genotyped as control, i.e., *TUB2*, *A. aurata* tubulin 2-coding gene; *GAPDH*, *A. aurata* glyceraldehyde 3-phosphate dehydrogenase-coding gene. Oligonucleotides are listed in **Supplementary Table 6**. Expected amplicon sizes were: *HPH*, 692 bp; *mRFP*, 591 bp; *FfERG27*, 648 bp; *TUB2*, 654 bp; *GAPDH*, 632 bp. P, positive control plasmids pTK144 (upper panel) and pTelFen (lower panel); H_2_O, no template negative control. Size marker, 1 kb plus (Invitrogen-Thermo Fisher, Waltham, MA, USA). (**D**) Confocal laser scanning microscopy of mycelium obtained from strain H1. Left, monomeric red fluorescence protein (mRFP) excitation at 561 nm and detection at 595-645 nm; middle, green fluorescence protein (GFP) excitation at 488 nm and detection at 505-555 nm (autofluorescence control); right, brightfield-mRFP channel overlay. Micrographs were taken with an SP8 confocal laser scanning microscope (Leica Microsystems GmbH, Wetzlar, Germany) using the LAS-X software. Scale bar: 25 µm.

## Discussion

The closest known extant relatives of the powdery mildew fungi belong to the *Arachnopezizaceae*, a family of wood- and litter-decaying saprotrophic fungi (Korf, 1951; Ekanayaka, 2019; Johnston *et al*., 2019; Vaghefi *et al*., 2022). In contrast to the obligate biotrophic powdery mildews, *Arachnopeziza* grows *in vitro* using common media and cultivation procedures (**Figure 1**; (Kosonen *et al*., 2020)), which implies that genetic modification and selection protocols may be feasible. In this work, we established high-quality genomic resources for the two species *A. aurata* and *A. aurelia*, including telomere-to-telomere genome assemblies (**Figure 2**) and coding gene and transposon annotations (**Table 1**, **Table 2**; **Supplementary Table 1** and **2**). We demonstrated that *A. aurata* is amenable to protoplast-based genetic modification by conferring transgene-mediated hygromycin resistance and red fluorescence (**Figure 5**, **Figure 6**), suggesting that *A. aurata* may represent a useful tool for the genetic study of the obligate biotrophic lifestyle in powdery mildew fungi. Fungi exhibit various lifestyles depending on the ecological niche they occupy (Lowe & Howlett, 2012). Saprotrophic fungi can decompose organic matter, including complex polysaccharides occurring in wood, bark, and plant litter. Obligate biotrophic fungi obtain carbohydrates and inorganic nutrients exclusively from a living host, while necrotrophic fungi kill and decompose their host while feeding on them. Obligate biotrophic plant pathogens, including powdery mildew fungi, rely on their host to obtain certain metabolites, as they have lost certain biosynthetic pathways (Spanu, 2012). Interestingly, saprotrophic fungi can undergo lifestyle transitions towards necrotrophy or facultative biotrophy under certain conditions and with compatible hosts (Seidl *et al*., 2015; Smith *et al*., 2017; Hill *et al*., 2022). The genomes of several obligate biotrophic plant pathogenic lineages, including powdery mildew fungi, rust fungi, and some oomycetes, are characterized by the loss of primary metabolic pathways such as thiamine biosynthesis and the inability to assimilate inorganic sulfur and nitrogen (Spanu *et al*., 2010; Baxter *et al*., 2010; Duplessis *et al*., 2011; Spanu, 2012). Since the *Arachnopezizaceae* are closely related to powdery mildew fungi, we queried *A. aurata* and *A. aurelia* for components of primary metabolism. Their primary metabolism profiles were more similar to that of the saprobic and necrotrophic fungus *B. cinerea*, including the presence of the thiamine biosynthesis pathway and sulfur and nitrogen assimilation pathways (**Figure 4A**). This is consistent with the ability of *A. aurata* and *A. aurelia* to grow under *in vitro* conditions (**Figure 1**). Further, obligate biotrophic plant pathogens like the powdery mildew fungi harbor genomes inflated in size due to the massive expansion of TEs, associated with the loss of the RIP pathway to control TE spread (Spanu *et al*., 2010; Kusch *et al*., 2024b). Both *A. aurata* and *A. aurelia* had average-sized genomes exhibiting TE contents below 5% (**Table 1**) and evidence of a functional RIP mechanism (**Figure 3**). Hence, the genome architectures of *A. aurata* and *A. aurelia* do not resemble those of obligate biotrophic plant pathogens like the powdery mildew fungi.

Saprobic fungi rely on a combination of secreted plant cell wall-degrading enzymes and the non-enzymatic Fenton reaction to decompose lignin and complex polysaccharides in wood and plant litter (Eastwood *et al*., 2011). Saprotrophic as well as necrotrophic fungi employ a wide array of CAZymes to degrade the complex polysaccharides derived from plant cell walls (Hage & Rosso, 2021), although the number of CAZymes can vary widely in ascomycetes, irrespective of their lifestyle (Zhao *et al*., 2013). In line with a saprotrophic lifestyle, and in contrast to the obligate biotrophic barley powdery mildew pathogen *B. hordei*, both *A. aurata* and *A. aurelia* have more than 500 proteins predicted to function as CAZymes (**Figure 4B**). Different from *B. hordei*, the CAZyme repertoire includes numerous glycosyl hydrolases and accessory enzymes (**Figure 4B**). This CAZyme profile resembles other saprotrophic fungi such as *Verticillium tricorpus* (Seidl *et al*., 2015). Taken together, *A. aurata* and *A. aurelia* exhibit the characteristic genomic signatures of a saprotrophic fungus able to decompose plant cell walls, and their genomes do not resemble the transposon-enriched architecture of the obligate biotrophic powdery mildew fungi.

Contrasting the genomic features expected of a saprotrophic fungus, we found more than 400 predicted effectors in each of the proteomes of *A. aurata* and *A. aurelia* (**Figure 4C**; **Table 2**). While functions related to carbohydrate modification and proteases were predicted for some of these candidate effectors (**Supplementary Table 1** and **2**), which is consistent with a saprobic lifestyle involving the decomposition of organic plant matter, around half of the proteins in both *Arachnopeziza* species had no predicted function and/or recognizable protein domain. However, the effector and CAZyme content of the genome alone may not always suffice to distinguish fungi of saprotrophic and pathogenic lifestyles. For instance, saprotrophic and plant-pathogenic *Fusarium* species exhibit a large overlap in effector and CAZyme repertoires, and lineages can be rather distinguished by a few lineage-specific effectors (Hill *et al*., 2022). In another example, the genome of the saprotrophic fungus *V. tricorpus* contains an effector repertoire similar to that of the vascular wilt pathogen *V. dahliae*, but an expanded set of CAZymes (Seidl *et al*., 2015). *V. tricorpus* is a saprotrophic fungus, but occasionally transitions to a pathogenic lifestyle (Nair *et al*., 2015; Farag *et al*., 2024). While such opportunistic infections by *Arachnopeziza* are not described, the large number of effectors in *A. aurata* and *A. aurelia* could facilitate pathogenic or mutualistic interactions with plants under certain conditions.

Several *Arachnopeziza* species seem to be closely associated with mosses such as *Sphagnum* and liverworts like *Ptilidium* (Stenroos *et al*., 2010; Kosonen *et al*., 2020). It is currently unclear if these associations represent close mutualistic or pathogenic interactions between *Arachnopeziza* species and mosses or liverworts. One possibility is that some of the potential effectors in *A. aurata* and *A. aurelia* serve to facilitate the interaction or communication with mosses. However, the species do not exhibit any of the auxotrophies that are typical for obligate biotrophic plant pathogens such as powdery mildews and rusts (**Figure 4A**). Likewise, *A. aurata* and *A. aurelia* do not seem to possess any polysaccharide lyases, which cleave uronic acid-containing polysaccharide chains (Lombard *et al*., 2010) and belong to a family of CAZymes often expanded in plant pathogens but absent in saprobic fungi (Karlsson *et al*., 2015). Therefore, such an interaction is likely to be either facultative or commensal. Indeed, saprotrophic fungi may harbor the capacity to enter into facultative biotrophic relationships with plant roots without causing disease symptoms (Smith *et al*., 2017), for instance by regulating their plant cell wall-degrading enzymes (Olson *et al*., 2012), which may otherwise cause activation of the plant immune system (Plett & Martin, 2011). A detailed study of the association between mosses and liverworts with *Arachnopeziza* is required to shed light on the quality of the interaction and has the potential to help better understand the saprobic and biotrophic lifestyles of these fungi.

Effectors can also mediate interactions with other microbes. For instance, the *V. dahliae* effectors VdAve1 and VdAMP2 modulate the microbiota composition within and outside the host plant, possibly to suppress harmful competitors or toxin-producing microbes (Snelders *et al*., 2020). Likewise, *A. aurelia* and *A. aurata* may employ effectors to help them compete or cooperate with other fungi and bacteria (Snelders *et al*., 2022).

Functional studies of powdery mildew effectors and other proteins are currently limited to indirect methods such as host-induced gene silencing (Nowara *et al*., 2010; Pedersen *et al*., 2012) and the screening of EMS- or UV-mutagenized populations (Barsoum *et al*., 2020; Bernasconi *et al*., 2024). Some attempts to genetically modify powdery mildew fungi in a targeted manner have been reported (Chaure *et al*., 2000; Martínez-Cruz *et al*., 2017), but a reliable and reproducible transformation system is lacking to date, possibly due to the obligate biotrophic lifestyle preventing *in vitro* cultivation. Here, we report that *A. aurata*, a saprotrophic fungus belonging to the family of the closest living relatives of powdery mildew fungi, can be genetically modified using a well-established protoplast-based transformation protocol (Leisen *et al*., 2020); **Figure 6**)). We argue that *A. aurata* holds great promise to study at least some of the powdery mildew proteins functionally, for instance concerning subcellular localization using fluorescent tags, protein-protein interactions, or post-translational modification profiles, all of which are currently not possible to assess in the fungus and can be only indirectly analyzed by *in planta* expression (e.g., (Pennington *et al*., 2019)). Further, targeted auxotrophies of, for instance, thiamine or the inability to assimilate sulfur or nitrogen could be introduced to study the genetic basis of obligate biotrophy in powdery mildew fungi.

## Methods

### Fungal strains and cultivation

The fungal strains *A. aurata* CBS 127674 and *A. aurelia* CBS 127675 were obtained in 2019 from the Westerdijk Fungal Biodiversity Institute at Utrecht University (CBS-KNAW, https://wi.knaw.nl/). Both fungi were routinely cultivated on maltose extract agar (MEA, 33.6 g L^-1^; Carl Roth GmbH, Karlsruhe, Germany) at 23 °C in the dark. Agar plugs with fungal mycelium were transferred to fresh MEA plates with a drop of 200 μL of sterile H_2_O every 2-4 weeks to maintain the original strains. Liquid cultures were grown in potato dextrose broth (PDB, 26.5 g L^-1^; Carl Roth GmbH) at 28 °C and 80 revolutions per minute (rpm) for 5-10 days. For testing different growth media, the fungi were further cultivated on potato dextrose agar (PDA, 39 g L^-1^; Sifin Diagnostics GmbH, Berlin, Germany), yeast peptone dextrose agar (YPDA, 65 g L^-1^; Carl Roth GmbH), or lysogeny broth agar (LB agar, containing 10 g L^-1^ tryptone, 5 g L^-1^ yeast extract, and 10 g L^-1^ sodium chloride; Carl Roth GmbH) at 23 °C for 24 days. For testing different growth temperatures, the fungi were grown on MEA at 16 °C, 23 °C, 28 °C, or 37 °C for 24 days.

### Antibiotic and fungicide susceptibility assays

*A. aurata* and *A. aurelia* were cultivated at 23 °C for 16 days on 1x MEA containing either of the following antibiotics or fungicides: hygromycin B (5-50 µg mL^-1^; Roche AG, Basel, Switzerland), fenhexamid (0.1-50 µg mL^-1^; Bayer AG, Leverkusen, Germany), geneticin G418 (5-50 µg mL^-1^; Santa Cruz Biotechnology, Dallas, USA), neomycin (5-50 µg mL^-1^; Duchefa Farma B. V., Haarlem, The Netherlands), kanamycin (5-400 µg mL^-1^; Carl Roth GmbH), streptomycin (5-400 µg mL^-1^; AppliChem GmbH, Darmstadt, Germany), ampicillin (5-400 µg mL^-1^; Duchefa Farma B. V.), and cefotaxime (5-400 µg mL^-1^; Carl Roth GmbH). Drug susceptibility was assessed by comparing mycelial growth at the varying concentrations of antibiotic or fungicide.

### Protoplast isolation

Fungal protoplasts were isolated based on a protocol for *Magnaporthe oryzae* (Leisen *et al*., 2020) with the following modifications: Mycelium balls were collected from liquid PDB cultures of *A. aurata* incubated at 80 rpm and 28 °C after 7-14 days, and mycelia were shredded using a sterile blender. The shredded hyphae were further incubated in PDB at 80 rpm and 28 °C for three days before transferring the mycelium to osmotic medium (1.2 M MgSO_4_, 10 mM NaPO_4_, pH 5.6 set with Na_2_HPO_4_) containing Glucanex (Novozymes A/S, Bagsværd, Denmark) for 3 h at 28 °C and 80 rpm. If protoplasts were detected via brightfield microscopy, they were filtered through sterile Miracloth (Merck KGaA, Darmstadt, Germany) and then washed using glacial TMS buffer (1 M sorbitol, 10 mM 3-(N-morpholino)propanesulfonic acid (MOPS), pH 6.3; Carl Roth GmbH). Then, protoplasts were centrifuged at 2000 x g and 4 °C for 15 min and resuspended in 1 mL of TMSC buffer (TMS with 46 mM CaCl_2_) and placed on ice; protoplasts were then ready for PEG-mediated transformation.

### PEG-mediated fungal transformation

The plasmids pTK144 (hygromycin resistance) and pTelFen (fenhexamid resistance) were used as transformation vectors (Leisen *et al*., 2020; Wegner *et al*., 2022). The DNA fragment containing the transgenes (*mRFP*, *HPH* and *FfERG27*, respectively) was amplified with the Phusion® High-Fidelity DNA Polymerase kit (New England Biolabs GmbH, Frankfurt a.M., Germany) using the oligonucleotides pTK144_FC_OE_Fw and pTK144_FC_OE_Rv (**Supplementary Table 3**) and the following thermal profile: initial denaturation at 98 °C for 30 s, 35 cycles of denaturation at 98 °C for 10 s, annealing at 60 °C for 30 s, and elongation at 72 °C for 2.5 min, and a final extension cycle at 72 °C for 5 min. PCR products were purified with the Monarch® Genomic DNA Purification Kit (New England Biolabs GmbH).

Protoplasts in TMSC were placed on ice for 10 min. Approximately 6 µg of the DNA was mixed with Tris-CaCl_2_ (10 mM Tris, 1 mM EDTA, 40 mM CaCl_2_, pH 6.3) to a final volume of 30-60 µL. 120 µL of protoplasts were added to the DNA, which then rested on ice for 10 min before adding 180 µL PEG solution (0.6 g mL^-1^ PEG3350 (Sigma-Aldrich, Munich, Germany), 1 M sorbitol, 10 mM MOPS, pH 6.3). Protoplasts were incubated at room temperature for 20 min. Then, protoplasts were regenerated in 50 mL of TB3 (0.2 g L^-1^ D(+)-saccharose, 3 mg L^-1^ yeast extract; Carl Roth GmbH) at 28 °C and 80 rpm for 3-5 days. Afterward, the fungal protoplasts were mixed with 50 mL of 2x MEA containing 10 µg mL^-1^ hygromycin B (Roche AG) or fenhexamid (Bayer AG) at 45 °C and then poured into a petri dish. The plates were incubated at 23 °C in the dark for 2-4 weeks. Colonies growing through the agar were transferred to new MEA plates containing hygromycin B or fenhexamid. Protoplasts without added DNA served as negative controls.

### Diagnostic polymerase chain reaction (PCR)

Diagnostic polymerase chain reaction (PCR) was conducted using isolated genomic DNA using the OneTaq® Quick-Load® DNA Polymerase system (New England Biolabs GmbH). PCRs were run with the following thermal profile: initial denaturation at 94 °C for 30 s, 35 cycles of denaturation at 94 °C for 30 s, annealing at 58-62 °C for 15-60 s, and elongation at 68 °C for 45 s, and a final extension cycle at 68 °C for 5 min. Oligonucleotides used in this work are listed in **Supplementary Table 6**. PCR products were assessed via electrophoresis at 80-100 V in a 1% agarose gel in 1x TAE buffer (40 mM Tris-HCl, 20 mM acetic acid, 1 mM EDTA, pH 8.0) using ethidium bromide (Carl Roth GmbH) as an intercalating DNA dye. The 6x TriTrack loading dye (NEB) was used as a DNA loading dye. DNA bands were visualized on a GelDocTM XR+ (Bio-Rad Laboratories GmbH, Feldkirchen, Germany).

### Whole transcriptome shotgun sequencing

*A. aurata* and *A. aurelia* were cultivated in PDB at 80 rpm and 28 °C for 7 days. The mycelia balls were flash-frozen in liquid N_2_ and crushed to a fine powder using a mortar and pestle, and RNA was isolated using the TRIzol® extraction protocol following the manufacturer’s instructions (Thermo Scientific, Karlsruhe, Germany). Genomic DNA removal was accomplished using RNase-free DNase I (Thermo Scientific). We determined the quality of RNA samples using microcapillary electrophoresis (2100 BioAnalyzier system; Agilent, Santa Clara, CA, USA) and RNA quantity by spectrophotometry (NanoDrop; Thermo Scientific) and spectrofluorimetry (Qubit; Thermo Scientific), confirming high-quality RNA for sequencing (RIN > 6.0, c[RNA] > 100 ng µL^−1^, m[RNA] > 1.5 µg). Stranded RNA sequencing with random oligomer priming, recovering total RNA and depletion of plant/animal ribosomal RNA were performed by the service provider Novogene Europe (Cambridge Science Park, UK), yielding 150-bp paired-end reads. The raw reads (available at NCBI/ENA/DDBJ at project accession PRJNA1128938) were trimmed with Trimmomatic v0.39 (Bolger *et al*., 2014) and quality control of the reads was conducted using FastQC v0.12.1 (Babraham Bioinformatics, Cambridge, UK).

### Whole genome sequencing and genome assembly

High molecular weight genomic DNA was obtained from *A. aurata* and *A. aurelia*, respectively, cultivated in PDB at 80 rpm and 28 °C for 7-14 days. Mycelia balls were flash-frozen in liquid N_2_ and crushed to a fine powder using a mortar and pestle. Then, DNA was isolated with a CTAB protocol according to (Feehan *et al*., 2017) with the modifications indicated in (Frantzeskakis *et al*., 2018). The DNA was further purified using the NucleoBond HMW DNA kit (Macherey-Nagel, Düren, Germany); DNA integrity was tested via a 0.6% agarose gel using ethidium bromide as an intercalating dye and run at 30 V for 3 h. DNA quantity was determined using the Qubit dsDNA-BR assay kit on a Qubit 4 (Thermo Fisher Scientific, Langerwehe, Germany).

DNA shotgun sequencing was performed using Illumina NovaSeq (NovaSeq 6000) technology with 1 µg input DNA at the service provider CeGaT (CeGaT, Tübingen, Germany), yielding 150-bp paired-end reads. We trimmed raw reads using Trimmomatic v0.39 (Bolger *et al*., 2014) and assessed read quality with FastQC v0.12.1 (Babraham Bioinformatics, Cambridge, UK). Long-read sequencing was performed by MinION (Oxford Nanopore Technologies, Oxford, US) with R9.4.1 flow cells and the Ligation Sequencing Kit SQK-LSK112; basecalling was done using guppy v0.15.3. All raw reads are available at NCBI/ENA/DDBJ at project accession PRJNA1128938.

We generated draft genome assemblies using the long reads with Canu v2.2 (Koren *et al*., 2017), Flye v2.9.2 (Kolmogorov *et al*., 2019) with options ‘`--iterations 3 –threads 12 --genome-size 42m --asm-coverage 50 -m 10000’`, and NextDenovo v2.5.2 (Hu *et al*., 2024) with configuration options ‘`sort_options = -m 10g -t 8`’ and ‘`nextgraph_options = -a 1 -q 10 -E 5000`’, and then merged the assemblies with quickmerge v0.3 (Chakraborty *et al*., 2016) to obtain the best draft assembly. We then remapped the Illumina short reads to the respective merged assemblies using the function ‘`bwa mem`‘ of BWA v0.7.17-r1188 (Li & Durbin, 2009) and polished the assembly using pilon v1.24 (Walker *et al*., 2014).

We obtained basic assembly statistics using Quast v5.2.0 (Gurevich *et al*., 2013) and assembly quality estimations with CRAQ v1.0.9 (Li *et al*., 2023). Further, we identified the 5.8S, 18S, 28S nuclear ribosomal DNA (nrDNA) and ITS sequences using the nrDNA sequences of *A. aurata* CBS127674 and *A. aurelia* CBS127675 from GenBank (accessions MH864617.1, MH876055.1, MH864618.1, and MH876056.1; alignments at **Supplementary Files 1** and **2**). Genome completeness was estimated using 1,706 ascomycete core genes from the ascomycota_odb10 database with compleasm v0.2.6 (Simão *et al*., 2015; Huang & Li, 2023).

### Telomere identification

Telomeres were manually identified at the ends of assembled contigs as telomeric repeats 5’-TTAGGG-3’ or 3’-CCCTAA-5’. To complete the sequence ends where telomeric repeats were not found, we used teloclip v0.0.4 (https://github.com/Adamtaranto/teloclip). Briefly, we mapped the long MinION nanopore reads to the respective assembly using Minimap2 v2.26-r1175 (Li, 2018) to retrieve nanopore reads at both ends of the assembly sequences containing telomeric repeats (5’-TTAGGG-3’) with options ‘`-k 20 -ax map-ont`’ and parsed the SAM files with SAMtools v1.18 (Li *et al*., 2009). Then, teloclip with options ‘`--motifs TTAGGG,TTAAGGG --matchAny`’ was used to filter reads mapping to chromosome ends, and with options ‘`--extractReads --extractDir SplitOverhang`’ to extract these reads. Then, the reads of each chromosome end were aligned via multiple sequence alignment using MAFFT v7.520 (Katoh & Standley, 2013) and Jalview v2.11.3.2 (Waterhouse *et al*., 2009) was employed to manually identify and extend scaffold ends via read alignment until the last aligning telomeric repeat, where available.

### Analysis of genome synteny

We used the functions nucmer for genome alignment and dnadiff, delta-filter with option ‘`-l 1000`’ and show-coords with ‘`-c -l -L 1000`’ for filtering from the MUMmer v4.4.0 package (Kurtz *et al*., 2004). We employed RIdeogram v0.2.2 (Hao et al., 2020) in R v4.3.1 (R Core Team, 2018) (www.r-project.org/) to visualize synteny, inversions, and translocations between the genome assemblies of *A. aurata* and *A. aurelia*.

### Genome map visualization

We visualized the genomic maps of *A. aurata* and *A. aurelia* as a circos diagram annotated with telomeres, centromeres, GC content, TE, and gene density. First, we used BEDtools v2.31.0 (Quinlan & Hall, 2010) to generate 5000-bp sliding windows with the function ‘makewindows’. Then, we calculated the GC content in each window with ‘`bedtools nuc`’ and further counted the number of annotated TEs and genes in each window via ‘`bedtools intersect`’. We searched for potential candidate centromeric regions by identifying windows covering between 30,000-150,000 bp where the gene content was zero or near zero and the TE content was at least 5 TEs per window. We then used circlize v0.4.10 (Gu *et al*., 2014) in R v4.3.1 (R Core Team, 2018) (www.r-project.org/) to visualize the genomic maps (see 03.circos_plot.md at https://github.com/stefankusch/arachnopeziza_analysis for a detailed script).

### Annotation of mitochondrial genomes

We used the online server of MFannot at https://megasun.bch.umontreal.ca/apps/mfannot/ accessed in 07/2024 (Lang *et al*., 2023) with the genetic code ‘4 Mold, Protozoan, and Coelenterate Mitochondria; Mycoplasma/Spiroplasma’ to discover and annotate mitochondrial genetic components. In addition, we used `blastn` and `tblastn` via NCBI BLAST+ v2.11.0 (Altschul *et al*., 1997) to search for coding genes and noncoding RNAs typically found in fungal mitochondrial genomes, i.e., ATP synthase subunit 6 (*atp6*, GenBank accession AGN49024.1), *atp8* (RKF65626.1), *atp9* (AGN49018.1), cytochrome c oxidase assembly factor 3 (*coa3-cc*, KAK6608516.1), mitochondrial cytochrome b (*cob*, ACL50595.1), cytochrome c oxidase subunit 1 (*cox1*, QPZ56210.1), (*cox2*, AGN49030.1), (*cox3*, AGN48999.1), NADH dehydrogenase subunit 1 (*nad1*, AGN49020.1), *nad2* (AGN49027.1), *nad3* (AGN49029.1), *nad4* (AGN49012.1), *nad4L* (AGN48994.1), *nad5* (AGN48996.1), *nad6* (AGN48998.1), mitochondrial small ribosomal subunit (*rns*,), mitochondrial large ribosomal subunit (*rnl*,), RNA component of ribonuclease P (*rnpB*, X93307.1), and ribosomal protein S3 (*rps3*, AGN49003.1).

### Annotation of transposable elements

We used the TEtrimmer pipeline (Qian *et al*., 2024) with the EDTA2-generated repeat libraries as input (Ou *et al*., 2019) to generate a consensus TE library for genome-wide annotation of TEs in the assemblies of *A. aurata* and *A. aurelia*. The two libraries were the user-defined libraries for genome-wide TE annotation with RepeatMasker v4.0.9 (http://www.repeatmasker.org) (Smit *et al*., 2016), yielding TE occupancy statistics, a TE annotation file, and the soft-masked genome assemblies. The sequence divergence of TE classes was calculated as described previously (Frantzeskakis *et al*., 2018, 2019).

### RIP analysis

Dinucleotide frequencies were calculated using RIPCAL v2.0 (Hane & Oliver, 2008), using FASTA sequences the whole genome assembly (without mitochondrial genome), coding sequences determined with BRAKER3, and repeat elements annotated with RepeatMasker (see above), i.e., *Ty1*/*Copia*, *Ty3*/*mdg4*, and DNA transposons as input. Then, the dinucleotide frequency indices were calculated in R v4.3.1 (R Core Team, 2018) (www.r-project.org/) (see 03.ripcal.md at https://github.com/stefankusch/arachnopeziza_analysis for a detailed script).

### Gene annotation

We used BRAKER3 v3.0.8 (Gabriel *et al*., 2021, 2023; Bruna *et al*., 2023) for evidence-based gene annotation of both *A. aurata* and *A. aurelia*. The respective RNA-seq data obtained in this work and the OrthoDB v11 Fungi protein dataset (https://www.orthodb.org/) (Kuznetsov *et al*., 2023) served as evidence datasets for BRAKER3 predictions. Reads were prepared for annotation by mapping to the respective genome assembly with HISAT2 (Kim *et al*., 2015) with ‘`--max-intronlen 1000 -k 10`’ and parsing the SAM files with SAMtools v1.18 (Li *et al*., 2009). We assessed the completeness of the gene annotations using BUSCO v5.5.0 (Simão *et al*., 2015) with ‘`-m protein`’ and the ascomycota_odb10 database.

We performed functional gene annotations using hmmscan from HMMer v3.4 (Potter *et al*., 2018) and InterProScan v5.73-104.0 (Jones *et al*., 2014) to predict functional protein domains, TMHMM v2.0c (Krogh *et al*., 2001) to detect putative transmembrane domains, SignalP5.0 (Almagro Armenteros *et al*., 2019) for secretion signals, EffectorP3.0 (Sperschneider & Dodds, 2021) for prediction of apoplastic/cytoplasmic effectors, dbCAN3 search at https://bcb.unl.edu/dbCAN2/ (accessed 07/2024) (Zheng *et al*., 2023) for the identification of candidate carbohydrate-active enzymes (CAZymes), antiSMASH v7.1.0 (Medema *et al*., 2011; Blin *et al*., 2023) to find genes encoding enzymes for secondary metabolic compounds, and primary metabolism components were annotated using GhostKOALA to identify components and KEGG Mapper to reconstruct pathways via https://www.kegg.jp/ (accessed 04/2025) (Kanehisa & Goto, 2000; Kanehisa *et al*., 2025). Orthologous proteins were identified using OrthoFinder v2.5.5 (Emms & Kelly, 2019).

Ribonuclease-like proteins in the proteomes of *A. aurata* and *A. aurelia* were analysed as follows: We extracted the sequences of putative secreted proteins containing PFAM domains (PF00445, PF06479) or InterPro domains (IPR036430, IPR016191) from the Ribonuclease T2-like superfamily. We used these sequences, i.e., AAURAT_002659-RB, AAURAT_005358-RA, AAURAT_010638-RA, and AAURAT_011677-RA for *A. aurata* and AAUREL_003350-RA, AAUREL_011122-RA, AAUREL_004672-RA for *A. aurelia*, as query for a protein BLAST (BLASTP) against the non-redundant protein database (nr on https://blast.ncbi.nlm.nih.gov/Blast.cgi accessed 04/2025) subset for a search within the Heliotales (taxonomy ID 5178) and randomly extracted protein sequences with high similarity to each of the queries. In addition, we included the sequences of three canonical RNase-like effector proteins, which were PM2 of *B. graminis* f.sp. *tritici* and CSEP0064 and CSEP0264 of *B. hordei* (Spanu, 2017). We conducted multiple sequence alignment using the MUSCLE algorithm and phylogenetic reconstruction using the function “build” of ETE3 3.1.3 (Huerta-Cepas *et al*., 2016) on the GenomeNet website (https://www.genome.jp/tools/ete/; accessed 04/2025). The maximum likelihood (ML) tree was inferred using RAxML v8.2.11 with model PROTGAMMAJTT and default parameters (Stamatakis, 2014).

### Data analysis and visualization

The R v4.3.1 software environment (R Core Team, 2018) (www.r-project.org/) was used for data analysis and plotting. Data analysis was facilitated by the packages tidyverse v2.0.0, dplyr v1.1.2, reshape2 v1.4.4, and scales v1.2.1. Bar plots and histograms were done using the R package ggplot2 v3.4.2 (Wickham, 2009). Synteny plots were generated with RIdeogram v0.2.2 (Hao et al., 2020) and circos plots with circlize v0.4.10 (Gu *et al*., 2014).

## Data availability statement

All raw RNA and DNA sequencing data generated in this study are deposited at https://www.ncbi.nlm.nih.gov/sra under BioProject ID PRJNA1128938. The draft genome assemblies for *Arachnopeziza aurata* CBS127674 and *A. aurelia* CBS127675 have been deposited at DDBJ/ENA/GenBank (accessions pending). Genome assemblies and gene annotations as used in this work are available at https://doi.org/10.5281/zenodo.15303401.

## Acknowledgments

We are grateful to Alex Wegener, Florencia Casanova, and Ulrich Schaffrath (RWTH Aachen University, Germany) for assistance with the PEG-mediated protoplast transformation protocol and for sharing the vectors pTK144 and pTelFen. The fungal strains were obtained from the Westerdijk Fungal Biodiversity Institute (CBS-KNAW), repository numbers CBS 127674 and CBS 127675. The analysis was performed with computing resources granted by RWTH Aachen University under project ID rwth0146.

## Author contributions

R.P., L.K., and S.K. designed the study; S.K. was responsible for experiment conception and planning. S.K. established the fungal *in vitro* cultures, supervised RNA and DNA extractions, and supervised cloning and genetic modification attempts. A.L. performed high molecular weight genomic DNA extraction and long-read DNA sequencing, established genetic modification protocols, and cloning and genotyping of strains. E.D. reproduced the genetic modification protocol, cloning, and genotyping. F.K. generated the samples for RNA-sequencing used for gene annotation. J.Q. conducted repeat masking and TE analysis of the *Arachnopeziza* genomes. H.I. did KEGG analysis. S.K. performed genome assemblies and polishing, annotation, comparative genomics, and data analysis. S.K. drafted figures and wrote the first draft of the manuscript, and S.K., L.K., and R.P. edited the manuscript. All authors read the manuscript and approved the final version.

## Abbreviations

bp: base pair
BLAST: Basic Local Alignment Search Tool
BUSCO: Benchmarking Universal Single-Copy Orthologs
CAZyme: carbohydrate-active enzyme
CDS: coding sequence
ITS: internal transcribed spacer
KEGG: Kyoto Encyclopedia of Genes and Genomes
LINE: long interspersed nuclear element
LTR: long terminal repeat
MEA: maltose extract agar
Mbp: million base pairs
nrDNA: nuclear ribosomal DNA
PCR: polymerase chain reaction
PDA: potato dextrose agar
PDB: potato dextrose broth
PEG: polyethylene glycol
PFAM: protein families
RFP: red fluorescent protein
RIP: repeat-induced point mutation
rpm: revolutions per minute
TE: transposable element

## Supplementary Materials

**Supplementary Table 1.** Functional predictions of *Arachnopeziza aurata* proteins annotated by BRAKER3.

**Supplementary Table 2**. Functional predictions of *Arachnopeziza aurelia* proteins annotated by BRAKER3.

**Supplementary Table 3**. KEGG pathway mapping summary of *Arachnopeziza aurata* and *A. aurelia* compared to *Botrytis cinerea* B05.10 and *Blumeria hordei* DH14.

**Supplementary Table 4**. Orthogroups of putative secreted proteins of *Arachnopeziza aurata*, *A. aurelia*, and *Blumeria hordei* DH14.

**Supplementary Table 5**. List of *Arachnopeziza* proteins containing ribonuclease-like domains.

**Supplementary Table 6**. Oligonucleotides used in this study.

**Supplementary File 1**. BLASTN search results for 5.8S, 18S, 28S nuclear ribosomal DNA (nrDNA) and ITS sequences (accessions MH864617.1, MH876055.1, MH864618.1, and MH876056.1) in the genome of *Arachnopeziza aurata*.

**Supplementary File 2**. BLASTN search results for 5.8S, 18S, 28S nuclear ribosomal DNA (nrDNA) and ITS sequences (accessions MH864617.1, MH876055.1, MH864618.1, and MH876056.1) in the genome of *Arachnopeziza aurelia*.

**Supplementary File 3**. TBLASTN search results for the repeat-induced point mutation (RIP) proteins *Masc1*, *Masc2*, *Rid-1*, and *Dim-2* (GenBank accession numbers AAC49849.1, AAC03766.1, XP_011392925.1, and XP_959891.1, respectively) in the genome of *Arachnopeziza aurata*.

**Supplementary File 4**. TBLASTN search results for the repeat-induced point mutation (RIP) proteins *Masc1*, *Masc2*, *Rid-1*, and *Dim-2* (GenBank accession numbers AAC49849.1, AAC03766.1, XP_011392925.1, and XP_959891.1, respectively) in the genome of *Arachnopeziza aurelia*.

**Supplementary File 5**. Multiple sequence alignment of *Arachnopeziza aurelia* AAUREL_003350-RA with different ribonucleases of the T1 family and ribonuclease-like effectors of plant pathogenic fungi.

**Supplementary File 6**. Multiple sequence alignment of protein sequences from *Arachnopeziza aurata* and *A. aurelia* containing ribonuclease-like domains, RNase-like effectors of *Blumeria hordei* and *B. graminis* f.sp. *tritici*, and ribonuclease-like proteins identified by a BLASTP search at https://blast.ncbi.nlm.nih.gov/Blast.cgi (accessed 04/2025).

## References

Aguileta G, De Vienne DM, Ross ON, Hood ME, Giraud T, Petit E, Gabaldón T. 2014. High Variability of Mitochondrial Gene Order among Fungi. Genome Biology and Evolution 6: 451–465.

Almagro Armenteros JJ, Tsirigos KD, Sønderby CK, Petersen TN, Winther O, Brunak S, von Heijne G, Nielsen H. 2019. SignalP 5.0 improves signal peptide predictions using deep neural networks. Nature Biotechnology 37: 420–423.

Altschul SF, Madden TL, Schäffer AA, Zhang J, Zhang Z, Miller W, Lipman DJ. 1997. Gapped BLAST and PSI-BLAST: a new generation of protein database search programs. Nucleic Acids Research 25: 3389–3402.

Baral HO. 2015. Nomenclatural novelties. Index Fungorum 225: 1–3.

Barsoum M, Kusch S, Frantzeskakis L, Schaffrath U, Panstruga R. 2020. Ultraviolet mutagenesis coupled with next-generation sequencing as a method for functional interrogation of powdery mildew genomes. Molecular Plant-Microbe Interactions 33: 1008–1021.

Baxter L, Tripathy S, Ishaque N, Boot N, Cabral A, Kemen E, Thines M, Ah-Fong A, Anderson R, Badejoko W, et al. 2010. Signatures of adaptation to obligate biotrophy in the *Hyaloperonospora arabidopsidis* genome. Science 330: 1549–1551.

Bernasconi Z, Stirnemann U, Heuberger M, Sotiropoulos AG, Graf J, Wicker T, Keller B, Sánchez-Martín J. 2024. Mutagenesis of wheat powdery mildew reveals a single gene controlling both NLR and tandem kinase-mediated immunity. Molecular Plant-Microbe Interactions 37: 264–276.

Blin K, Shaw S, Augustijn HE, Reitz ZL, Biermann F, Alanjary M, Fetter A, Terlouw BR, Metcalf WW, Helfrich EJN, et al. 2023. antiSMASH 7.0: New and improved predictions for detection, regulation, chemical structures and visualisation. Nucleic Acids Research 51: W46–W50.

Bolger AM, Lohse M, Usadel B. 2014. Trimmomatic: A flexible trimmer for Illumina sequence data. Bioinformatics 30: 2114–2120.

Braun U, Cook RTA. 2012. Taxonomic Manual of the Erysiphales (Powdery Mildews). Utrecht, The Netherlands: CBS-KNAW Fungal Biodiversity Centre.

Bruna T, Lomsadze A, Borodovsky M. 2023. A new gene finding tool GeneMark-ETP significantly improves the accuracy of automatic annotation of large eukaryotic genomes.

Chakraborty M, Baldwin-Brown JG, Long AD, Emerson JJ. 2016. Contiguous and accurate *de novo* assembly of metazoan genomes with modest long read coverage. Nucleic Acids Research 44: e147.

Chaure P, Gurr SJ, Spanu P. 2000. Stable transformation of *Erysiphe graminis* an obligate biotrophic pathogen of barley. Nature Biotechnology 18: 205–207.

Cohrs KC, Burbank J, Schumacher J. 2017. A new transformant selection system for the gray mold fungus *Botrytis cinerea* based on the expression of fenhexamid-insensitive ERG27 variants. Fungal Genetics and Biology 100: 42–51.

Credle JJ, Finer-Moore JS, Papa FR, Stroud RM, Walter P. 2005. On the mechanism of sensing unfolded protein in the endoplasmic reticulum. Proceedings of the National Academy of Sciences 102: 18773–18784.

Duplessis S, Cuomo CA, Lin Y-C, Aerts A, Tisserant E, Veneault-Fourrey C, Joly DL, Hacquard S, Amselem J, Cantarel BL, et al. 2011. Obligate biotrophy features unraveled by the genomic analysis of rust fungi. Proceedings of the National Academy of Sciences 108: 9166–9171.

Eastwood DC, Floudas D, Binder M, Majcherczyk A, Schneider P, Aerts A, Asiegbu FO, Baker SE, Barry K, Bendiksby M, et al. 2011. The plant cell wall–decomposing machinery underlies the functional diversity of forest fungi. Science 333: 762–765.

Ekanayaka A. 2019. Preliminary classification of Leotiomycetes. Mycosphere 10: 310–489.

Emms DM, Kelly S. 2019. OrthoFinder: phylogenetic orthology inference for comparative genomics. Genome Biology 20: 238.

Farag FM, Arafa RA, Abou-Zeid MA, Aloufi AS, Abd El Moneim D, Ghebrial EWR. 2024. First record of *Verticillium tricorpus* as a causal agent of *Verticillium* wilt disease in Okra. *Journal of Plant Diseases and Protection* 131: 557–569.

Feehan JM, Scheibel KE, Bourras S, Underwood W, Keller B, Somerville SC. 2017. Purification of high molecular weight genomic DNA from powdery mildew for long-read sequencing. Journal of Visualized Experiments: JoVE: e55463.

Franco MEE, López SMY, Medina R, Lucentini CG, Troncozo MI, Pastorino GN, Saparrat MCN, Balatti PA. 2017. The mitochondrial genome of the plant-pathogenic fungus *Stemphylium lycopersici* uncovers a dynamic structure due to repetitive and mobile elements (M-J Virolle, Ed.). PLOS ONE 12: e0185545.

Frantzeskakis L, Kracher B, Kusch S, Yoshikawa-Maekawa M, Bauer S, Pedersen C, Spanu PD, Maekawa T, Schulze-Lefert P, Panstruga R. 2018. Signatures of host specialization and a recent transposable element burst in the dynamic one-speed genome of the fungal barley powdery mildew pathogen. BMC Genomics 19: 381.

Frantzeskakis L, Németh MZ, Barsoum M, Kusch S, Kiss L, Takamatsu S, Panstruga R. 2019. The *Parauncinula polyspora* draft genome provides insights into patterns of gene erosion and genome expansion in powdery mildew fungi. mBio 10: 381.

Fuckel KWGL. 1870. Symbolae mycologicae. Beiträge zur Kenntniss der rheinischen Pilze. Jahrbücher des Nassauischen Vereins für Naturkunde 23–24: 459.

Gabriel L, Brůna T, Hoff KJ, Ebel M, Lomsadze A, Borodovsky M, Stanke M. 2023. BRAKER3: Fully automated genome annotation using RNA-seq and protein evidence with GeneMark-ETP, AUGUSTUS and TSEBRA.

Gabriel L, Hoff KJ, Brůna T, Borodovsky M, Stanke M. 2021. TSEBRA: Transcript selector for BRAKER. BMC Bioinformatics 22: 566.

Gladyshev E. 2017. Repeat-induced point mutation and other genome defense mechanisms in fungi (J Heitman and EH Stukenbrock, Eds.). Microbiology Spectrum 5: 5.4.02.

Glawe DA. 2008. The powdery mildews: A review of the world’s most familiar (yet poorly known) plant pathogens. Annual Review of Phytopathology 46: 27–51.

Gu Z, Gu L, Eils R, Schlesner M, Brors B. 2014. *circlize* implements and enhances circular visualization in R. Bioinformatics 30: 2811–2812.

Gurevich A, Saveliev V, Vyahhi N, Tesler G. 2013. QUAST: Quality assessment tool for genome assemblies. Bioinformatics 29: 1072–1075.

Hage H, Rosso M-N. 2021. Evolution of fungal carbohydrate-active enzyme portfolios and adaptation to plant cell-wall polymers. Journal of Fungi 7: 185.

Han J-G, Hosoya T, Sung G-H, Shin H-D. 2014. Phylogenetic reassessment of *Hyaloscyphaceae* sensu lato (Helotiales, Leotiomycetes) based on multigene analyses. Fungal Biology 118: 150–167.

Hane JK, Oliver RP. 2008. RIPCAL: A tool for alignment-based analysis of repeat-induced point mutations in fungal genomic sequences. BMC Bioinformatics 9: 478.

Hao Z, Lv D, Ge Y, Shi J, Weijers D, Yu G, Chen J. 2020. *RIdeogram* : Drawing SVG graphics to visualize and map genome-wide data on the idiograms. PeerJ Computer Science 6: e251.

Hill R, Buggs RJA, Vu DT, Gaya E. 2022. Lifestyle transitions in fusarioid fungi are frequent and lack clear genomic signatures. Molecular Biology and Evolution 39: msac085.

Hu J, Wang Z, Sun Z, Hu B, Ayoola AO, Liang F, Li J, Sandoval JR, Cooper DN, Ye K, et al. 2024. NextDenovo: An efficient error correction and accurate assembly tool for noisy long reads. Genome Biology 25: 107.

Huang N, Li H. 2023. compleasm: A faster and more accurate reimplementation of BUSCO (T Marschall, Ed.). Bioinformatics 39: btad595.

Huerta-Cepas J, Serra F, Bork P. 2016. ETE 3: Reconstruction, analysis, and visualization of phylogenomic data. Molecular Biology and Evolution 33: 1635–1638.

James TY, Pelin A, Bonen L, Ahrendt S, Sain D, Corradi N, Stajich JE. 2013. Shared signatures of parasitism and phylogenomics unite Cryptomycota and Microsporidia. Current Biology 23: 1548– 1553.

Jia J, Xue Q. 2009. Codon usage biases of transposable elements and host nuclear genes in *Arabidopsis Thaliana* and *Oryza Sativa*. Genomics, Proteomics & Bioinformatics 7: 175–184.

Johnston PR, Quijada L, Smith CA, Baral H-O, Hosoya T, Baschien C, Pärtel K, Zhuang W-Y, Haelewaters D, Park D, et al. 2019. A multigene phylogeny toward a new phylogenetic classification of *Leotiomycetes*. IMA Fungus 10.

Jones P, Binns D, Chang H-Y, Fraser M, Li W, McAnulla C, McWilliam H, Maslen J, Mitchell A, Nuka G, et al. 2014. InterProScan 5: Genome-scale protein function classification. Bioinformatics 30: 1236–1240.

Kanehisa M, Furumichi M, Sato Y, Matsuura Y, Ishiguro-Watanabe M. 2025. KEGG: Biological systems database as a model of the real world. Nucleic Acids Research 53: D672–D677.

Kanehisa M, Goto S. 2000. KEGG: Kyoto Encyclopedia of Genes and Genomes. Nucleic Acids Research 28: 27–30.

Karlsson M, Durling MB, Choi J, Kosawang C, Lackner G, Tzelepis GD, Nygren K, Dubey MK, Kamou N, Levasseur A, et al. 2015. Insights on the evolution of mycoparasitism from the genome of *Clonostachys rosea*. Genome Biology and Evolution 7: 465–480.

Katoh K, Standley DM. 2013. MAFFT multiple sequence alignment software version 7: Improvements in performance and usability. Molecular Biology and Evolution 30: 772–780.

Keller NP. 2019. Fungal secondary metabolism: Regulation, function and drug discovery. Nature Reviews Microbiology 17: 167–180.

Kemen AC, Agler MT, Kemen E. 2015. Host–microbe and microbe–microbe interactions in the evolution of obligate plant parasitism. New Phytologist 206: 1207–1228.

Kim D, Langmead B, Salzberg SL. 2015. HISAT: A fast spliced aligner with low memory requirements. Nature Methods 12: 357–360.

Kirk PM, Cannon PF, Stalpers JA. 2008. Dictionary of the fungi. Wallingford, Oxon, UK: CABI.

Kiss L, Vaghefi N, Bransgrove K, Dearnaley JDW, Takamatsu S, Tan YP, Marston C, Liu S-Y, Jin D-N, Adorada DL, et al. 2020. Australia: A continent without native powdery mildews? The first comprehensive catalog indicates recent introductions and multiple host range expansion events, and leads to the re-discovery of *Salmonomyces* as a new lineage of the Erysiphales. Frontiers in Microbiology 11: 1571.

Kolmogorov M, Yuan J, Lin Y, Pevzner PA. 2019. Assembly of long, error-prone reads using repeat graphs. Nature Biotechnology 37: 540–546.

Koren S, Walenz BP, Berlin K, Miller JR, Bergman NH, Phillippy AM. 2017. Canu: scalable and accurate long-read assembly via adaptive k-mer weighting and repeat separation. Genome Research 27: 722–736.

Korf RP. 1951. A monograph of the *Arachnopezizeae*. Lloydia 14: 129–180.

Kosonen T, Huhtinen S, Hansen K. 2020. Taxonomy and systematics of *Hyaloscyphaceae* and *Arachnopezizaceae*. Persoonia - Molecular Phylogeny and Evolution of Fungi 46: 26–62.

Krogh A, Larsson B, von Heijne G, Sonnhammer ELL. 2001. Predicting transmembrane protein topology with a hidden markov model: Application to complete genomes. Journal of Molecular Biology 305: 567–580.

Kubicek CP, Starr TL, Glass NL. 2014. Plant cell wall-degrading enzymes and their secretion in plant-pathogenic fungi. Annual Review of Phytopathology 52: 427–451.

Kurtz S, Phillippy A, Delcher AL, Smoot M, Shumway M, Antonescu CM, Salzberg SL. 2004. Versatile and open software for comparing large genomes. Genome Biology 5: R12.

Kusch S, Frantzeskakis L, Lassen BD, Kümmel F, Pesch L, Barsoum M, Walden KD, Panstruga R. 2024a. A fungal plant pathogen overcomes *mlo*-mediated broad-spectrum disease resistance by rapid gene loss. New Phytologist 244: 962–979.

Kusch S, Qian J, Loos A, Kümmel F, Spanu PD, Panstruga R. 2024b. Long-term and rapid evolution in powdery mildew fungi. Molecular Ecology 33: e16909.

Kuznetsov D, Tegenfeldt F, Manni M, Seppey M, Berkeley M, Kriventseva EV, Zdobnov EM. 2023. OrthoDB v11: Annotation of orthologs in the widest sampling of organismal diversity. Nucleic Acids Research 51: D445–D451.

Lang BF, Beck N, Prince S, Sarrasin M, Rioux P, Burger G. 2023. Mitochondrial genome annotation with MFannot: A critical analysis of gene identification and gene model prediction. Frontiers in Plant Science 14: 1222186.

Leisen T, Bietz F, Werner J, Wegner A, Schaffrath U, Scheuring D, Willmund F, Mosbach A, Scalliet G, Hahn M. 2020. CRISPR/Cas with ribonucleoprotein complexes and transiently selected telomere vectors allows highly efficient marker-free and multiple genome editing in Botrytis cinerea (M Bolton, Ed.). PLOS Pathogens 16: e1008326.

Li H. 2018. Minimap2: Pairwise alignment for nucleotide sequences. Bioinformatics 34: 3094–3100.

Li H, Durbin R. 2009. Fast and accurate short read alignment with Burrows–Wheeler transform. Bioinformatics 25: 1754–1760.

Li H, Handsaker B, Wysoker A, Fennell T, Ruan J, Homer N, Marth G, Abecasis G, Durbin R. 2009. The Sequence Alignment/Map format and SAMtools. Bioinformatics 25: 2078–2079.

Li K, Xu P, Wang J, Yi X, Jiao Y. 2023. Identification of errors in draft genome assemblies at single-nucleotide resolution for quality assessment and improvement. Nature Communications 14: 6556.

Lichius A, Ruiz DM, Zeilinger S. 2020. Genetic Transformation of Filamentous Fungi: Achievements and Challenges. In: Nevalainen H, ed. Grand Challenges in Biology and Biotechnology. Grand Challenges in Fungal Biotechnology. Cham: Springer International Publishing, 123–164.

Liu W, Cai Y, Zhang Q, Shu F, Chen L, Ma X, Bian Y. 2020. Subchromosome-scale nuclear and complete mitochondrial genome characteristics of *Morchella crassipes*. International Journal of Molecular Sciences 21: 483.

Lombard V, Bernard T, Rancurel C, Brumer H, Coutinho PM, Henrissat B. 2010. A hierarchical classification of polysaccharide lyases for glycogenomics. Biochemical Journal 432: 437–444.

Lowe RGT, Howlett BJ. 2012. Indifferent, affectionate, or deceitful: Lifestyles and secretomes of fungi (J Heitman, Ed.). PLoS Pathogens 8: e1002515.

Margolin BS, Garrett-Engele PW, Stevens JN, Fritz DY, Garrett-Engele C, Metzenberg RL, Selker EU. 1998. A methylated *Neurospora* 5S rRNA pseudogene contains a transposable element inactivated by repeat-induced point mutation. Genetics 149: 1787–1797.

Martínez-Cruz J, Romero D, de Vicente A, Pérez-García A. 2017. Transformation of the cucurbit powdery mildew pathogen *Podosphaera xanthii* by *Agrobacterium tumefaciens*. New Phytologist 213: 1961–1973.

Medema MH, Blin K, Cimermancic P, de Jager V, Zakrzewski P, Fischbach MA, Weber T, Takano E, Breitling R. 2011. antiSMASH: Rapid identification, annotation and analysis of secondary metabolite biosynthesis gene clusters in bacterial and fungal genome sequences. Nucleic Acids Research 39: W339–W346.

Mohanta TK, Bae H. 2015. The diversity of fungal genome. Biological Procedures Online 17: 8.

Müller MC, Praz CR, Sotiropoulos AG, Menardo F, Kunz L, Schudel S, Oberhänsli S, Poretti M, Wehrli A, Bourras S, et al. 2019. A chromosome-scale genome assembly reveals a highly dynamic effector repertoire of wheat powdery mildew. New Phytologist 221: 2176–2189.

Nair PVR, Wiechel TJ, Crump NS, Taylor PWJ. 2015. First report of *Verticillium tricorpus* causing *Verticillium* wilt in potatoes in Australia. Plant Disease 99: 731–731.

Nguyen TAN, Higa T, Shiina A, Utami YD, Hiruma K. 2023. Exploring the roles of fungal-derived secondary metabolites in plant-fungal interactions. Physiological and Molecular Plant Pathology 125: 102021.

Nowara D, Gay A, Lacomme C, Shaw J, Ridout CJ, Douchkov D, Hensel G, Kumlehn J, Schweizer P. 2010. HIGS: host-induced gene silencing in the obligate biotrophic fungal pathogen *Blumeria graminis*. The Plant Cell 22: 3130–3141.

Oggenfuss U, Croll D. 2023. Recent transposable element bursts are associated with the proximity to genes in a fungal plant pathogen (B Thomma, Ed.). PLOS Pathogens 19: e1011130.

Olson Å, Aerts A, Asiegbu F, Belbahri L, Bouzid O, Broberg A, Canbäck B, Coutinho PM, Cullen D, Dalman K, et al. 2012. Insight into trade-off between wood decay and parasitism from the genome of a fungal forest pathogen. New Phytologist 194: 1001–1013.

Ou S, Su W, Liao Y, Chougule K, Agda JRA, Hellinga AJ, Lugo CSB, Elliott TA, Ware D, Peterson T, et al. 2019. Benchmarking transposable element annotation methods for creation of a streamlined, comprehensive pipeline. Genome Biology 20: 275.

Pedersen C, van Themaat EVL, McGuffin LJ, Abbott JC, Burgis TA, Barton G, Bindschedler LV, Lu X, Maekawa T, Weßling R, et al. 2012. Structure and evolution of barley powdery mildew effector candidates. BMC Genomics 13: 694.

Pennington HG, Jones R, Kwon S, Bonciani G, Thieron H, Chandler T, Luong P, Morgan SN, Przydacz M, Bozkurt T, et al. 2019. The fungal ribonuclease-like effector protein CSEP0064/BEC1054 represses plant immunity and interferes with degradation of host ribosomal RNA (PN Dodds, Ed.). PLoS Pathogens 15: e1007620.

Penselin D, Münsterkötter M, Kirsten S, Felder M, Taudien S, Platzer M, Ashelford K, Paskiewicz KH, Harrison RJ, Hughes DJ, et al. 2016. Comparative genomics to explore phylogenetic relationship, cryptic sexual potential and host specificity of Rhynchosporium species on grasses. BMC Genomics 17: 953.

Plett JM, Martin F. 2011. Blurred boundaries: Lifestyle lessons from ectomycorrhizal fungal genomes. Trends in Genetics 27: 14–22.

Potter SC, Luciani A, Eddy SR, Park Y, Lopez R, Finn RD. 2018. HMMER web server: 2018 update. Nucleic Acids Research 46: W200–W204.

Qian J, Xue H, Ou S, Storer J, Fürtauer L, Wildermuth MC, Kusch S, Panstruga R. 2024. TEtrimmer: A novel tool to automate the manual curation of transposable elements.

Quinlan AR, Hall IM. 2010. BEDTools: A flexible suite of utilities for comparing genomic features. Bioinformatics 26: 841–842.

R Core Team. 2018. R: A language and environment for statistical computing. R Foundation for Statistical Computing, Vienna, Austria.

Robey MT, Caesar LK, Drott MT, Keller NP, Kelleher NL. 2021. An interpreted atlas of biosynthetic gene clusters from 1,000 fungal genomes. Proceedings of the National Academy of Sciences 118: e2020230118.

Sabelleck B, Deb S, Levecque SCJ, Freh M, Reinstädler A, Spanu PD, Thordal-Christensen H, Panstruga R. 2025. A powdery mildew core effector protein targets the host endosome tethering complexes HOPS and CORVET in barley. Plant Physiology: kiaf067.

Schwarzbach E. 1979. Response to selection for virulence against the *ml-o* based mildew resistance in barley, not fitting the gene-for-gene hypothesis. Barley Genetics Newsletter 9: 85–89.

Seidl MF, Faino L, Shi-Kunne X, Van Den Berg GCM, Bolton MD, Thomma BPHJ. 2015. The genome of the saprophytic fungus *Verticillium tricorpus* reveals a complex effector repertoire resembling that of its pathogenic relatives. Molecular Plant-Microbe Interactions® 28: 362–373.

Seif E, Leigh J, Liu Y, Roewer I, Forget L, Lang BF. 2005. Comparative mitochondrial genomics in zygomycetes: Bacteria-like RNase P RNAs, mobile elements and a close source of the group I intron invasion in angiosperms. Nucleic Acids Research 33: 734–744.

Simão FA, Waterhouse RM, Ioannidis P, Kriventseva EV, Zdobnov EM. 2015. BUSCO: assessing genome assembly and annotation completeness with single-copy orthologs. *Bioinformatics (Oxford*, England) 31: 3210–3212.

Smit AFA, Hubley R, Green P. 2016. Masker Open-4.0. 2013–2015.

Smith GR, Finlay RD, Stenlid J, Vasaitis R, Menkis A. 2017. Growing evidence for facultative biotrophy in saprotrophic fungi: Data from microcosm tests with 201 species of wood-decay basidiomycetes. New Phytologist 215: 747–755.

Snelders NC, Rovenich H, Petti GC, Rocafort M, van den Berg GCM, Vorholt JA, Mesters JR, Seidl MF, Nijland R, Thomma BPHJ. 2020. Microbiome manipulation by a soil-borne fungal plant pathogen using effector proteins. Nature Plants 6: 1365–1374.

Snelders NC, Rovenich H, Thomma BPHJ. 2022. Microbiota manipulation through the secretion of effector proteins is fundamental to the wealth of lifestyles in the fungal kingdom. FEMS Microbiology Reviews 46: fuac022.

Spanu PD. 2012. The genomics of obligate (and nonobligate) biotrophs. Annual Review of Phytopathology 50: 91–109.

Spanu PD. 2017. Cereal immunity against powdery mildews targets RNase-Like Proteins associated with Haustoria (RALPH) effectors evolved from a common ancestral gene. New Phytologist 213: 969– 971.

Spanu PD, Abbott JC, Amselem J, Burgis TA, Soanes DM, Stüber K, van Themaat EVL, Brown JKM, Butcher SA, Gurr SJ, et al. 2010. Genome expansion and gene loss in powdery mildew fungi reveal tradeoffs in extreme parasitism. *Science (New York*, N.Y.) 330: 1543–1546.

Spanu PD, Panstruga R. 2017. Editorial: Biotrophic plant-microbe interactions. Frontiers in Plant Science 8: 192.

Sperschneider J, Dodds P. 2021. EffectorP 3.0: Prediction of apoplastic and cytoplasmic effectors in fungi and oomycetes. Molecular Plant-Microbe Interactions 35: 146–156.

Stamatakis A. 2014. RAxML version 8: A tool for phylogenetic analysis and post-analysis of large phylogenies. Bioinformatics 30: 1312–1313.

Stenroos S, Laukka T, Huhtinen S, Döbbeler P, Myllys L, Syrjänen K, Hyvönen J. 2010. Multiple origins of symbioses between ascomycetes and bryophytes suggested by a five-gene phylogeny. Cladistics 26: 281–300.

Vaghefi N, Kusch S, Nemeth MZ, Seress D, Braun U, Takamatsu S, Panstruga R, Kiss L. 2022. Beyond nuclear ribosomal DNA sequences: Evolution, taxonomy, and closest known saprobic relatives of powdery mildew fungi (*Erysiphaceae*) inferred from their first comprehensive genome-scale phylogenetic analyses. Frontiers in Microbiology 13: 903024.

Van Kan JAL, Stassen JHM, Mosbach A, Van Der Lee TAJ, Faino L, Farmer AD, Papasotiriou DG, Zhou S, Seidl MF, Cottam E, et al. 2017. A gapless genome sequence of the fungus *Botrytis cinerea*. Molecular Plant Pathology 18: 75–89.

Walker BJ, Abeel T, Shea T, Priest M, Abouelliel A, Sakthikumar S, Cuomo CA, Zeng Q, Wortman JR, Young SK, et al. 2014. Pilon: An integrated tool for comprehensive microbial variant detection and genome assembly improvement (J Wang, Ed.). PLoS ONE 9: e112963.

Waterhouse AM, Procter JB, Martin DMA, Clamp M, Barton GJ. 2009. Jalview Version 2 - a multiple sequence alignment editor and analysis workbench. Bioinformatics 25: 1189–1191.

Wegner A, Wirtz L, Leisen T, Hahn M, Schaffrath U. 2022. Fenhexamid - an efficient and inexpensive fungicide for selection of *Magnaporthe oryzae* transformants. European Journal of Plant Pathology 162: 697–707.

Wickham H. 2009. ggplot2: Elegant Graphics for Data Analysis (R Gentleman, K Hornik, and G Parmigiani, Eds.). New York: Springer-Verlag.

Zaccaron AZ, De Souza JT, Stergiopoulos I. 2021. The mitochondrial genome of the grape powdery mildew pathogen *Erysiphe necator* is intron rich and exhibits a distinct gene organization. Scientific Reports 11: 13924.

Zaccaron AZ, Stergiopoulos I. 2021. Characterization of the mitochondrial genomes of three powdery mildew pathogens reveals remarkable variation in size and nucleotide composition. Microbial Genomics 7: 000720.

Zhao Z, Liu H, Wang C, Xu J-R. 2013. Comparative analysis of fungal genomes reveals different plant cell wall degrading capacity in fungi. BMC Genomics 14: 274.

Zheng J, Ge Q, Yan Y, Zhang X, Huang L, Yin Y. 2023. dbCAN3: Automated carbohydrate-active enzyme and substrate annotation. Nucleic Acids Research 51: W115–W121.

