## Supplementary Figures for "Resources for molecular studies of unculturable obligate biotrophic fungal plant pathogens using their saprotrophic relatives"

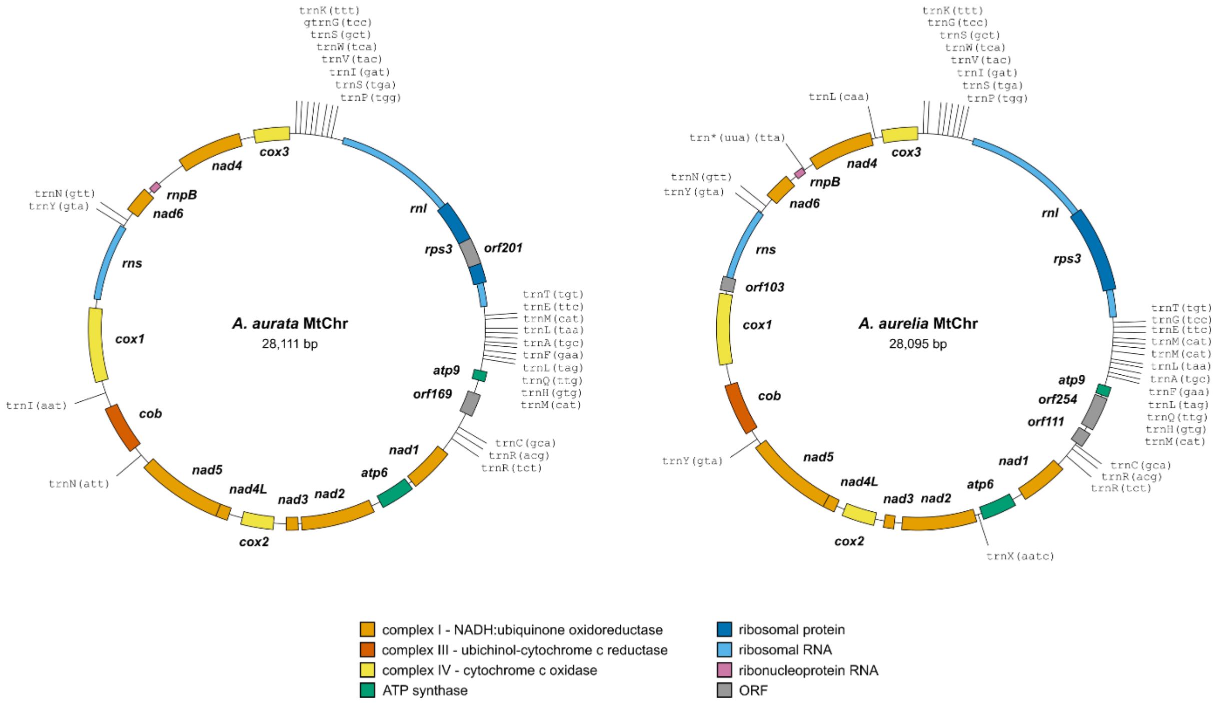


**Supplementary Figure 1. The mitochondrial genomes of *A. aurata* and *A. aurelia* are compact and seem to lack the ATP synthase component *atp8*.** The circular mitochondrial genome assemblies were recovered from the initial whole genome assemblies of *A. aurata* CBS127647 (left) and *A. aurelia* CBS127675 (right). The mitochondrial genes were annotated using MFannot (Lang *et al.*, 2023) and additional BLASTN and TBLASTN searches to recover missing components. The thick bars indicate the coding genes for the mitochondrial respiratory machinery color-coded by the mitochondrial respiratory complex, i.e., NADH:ubiquinone oxidoreductase (complex I, orange), ubiquinol-cytochrome c oxidoreductase (complex III or $bc_1$ complex, dark orange), cytochrome c oxidase (complex IV, yellow), ATP synthase (green), and ribosomal protein *rps3* (dark blue); other open reading frames are shown in grey. Thin bars indicate noncoding RNAs (*rns*, small ribosomal RNA, light blue; *rnl*, large ribosomal RNA, light blue; *rnpB*, mitochondrial RNaseP-RNA, purple). The mitochondrial tRNAs are added as labels.

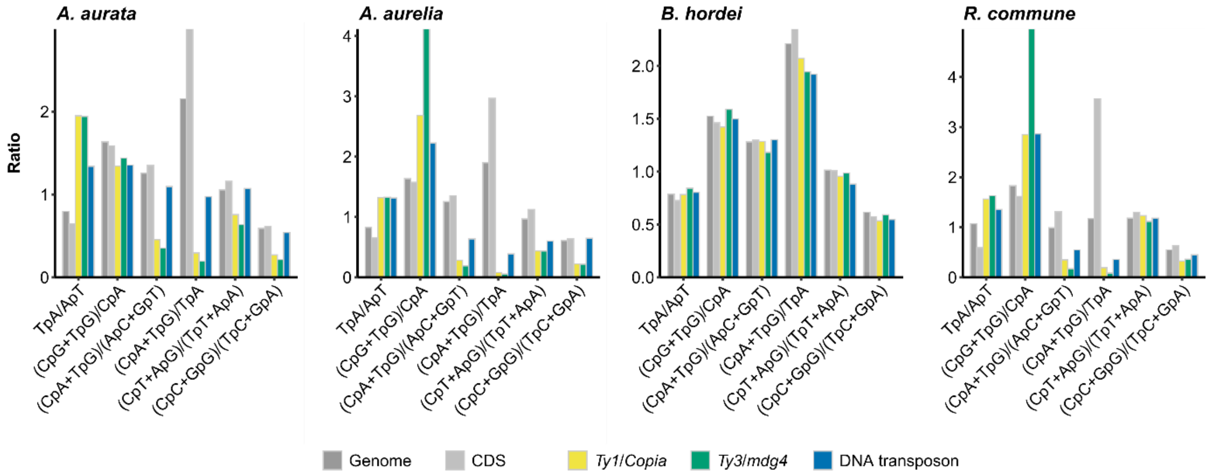


**Supplementary Figure 2. The genomes of *Arachnopeziza aurata* and *A. aurelia* show signs of TE degeneration via RIP.** We performed RIP index analysis based on dinucleotide ratios via RIPCAL (Hane & Oliver, 2008) on the genome assemblies of *A. aurata* CBS127674 and *A. aurelia* CBS127675 with the assemblies of the barley powdery mildew *Blumeria hordei* DH14 (Frantzeskakis *et al.*, 2018) and the barley leaf scald disease pathogen *Rhynchosporium commune* UK7 (Penselin *et al.*, 2016). Panels are displayed in this order from left to right. The bar graphs show the ratios of repetitive sequence dinucleotide analysis genome-wide (dark grey), for coding sequences (CDS; light grey), *Ty1*/*Copia* LTR elements (yellow), *Ty3*/*mdg4* LTR elements (orange), and DNA transposons (blue). The respective RIP index represented by dinucleotide ratios is indicated on the x-axis, and the y-axis displays the respective ratio.

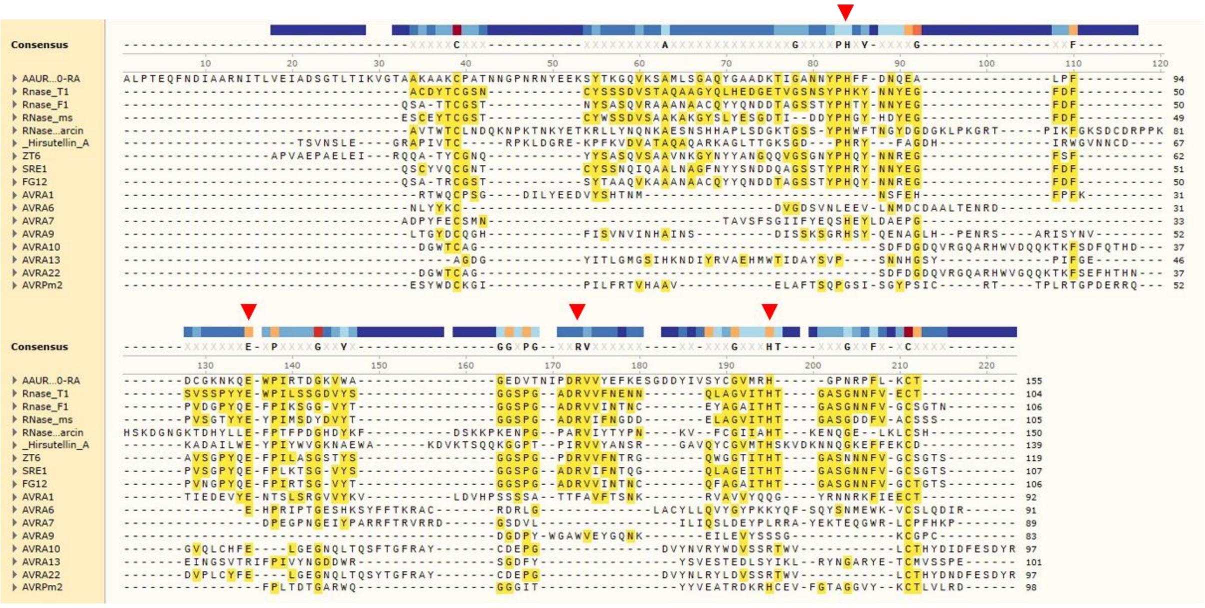


**Supplementary Figure 3. Alignment of AAUREL_003350-RA with different ribonucleases of the T1 family and ribonuclease-like effectors of plant-pathogenic fungi.** The multiple sequence alignment of the amino acid sequences was generated with SnapGene v6.0.2 and the implemented MAFFT v7.471 aligner (SnapGene® software (from Dotmatics; available at snapgene.com)). The residues required for guanyl-specific ribonuclease activity of full-length RNase T1 are H66, E84, R103, H118, which correspond to P40, E58, R77, H92, respectively (numbering without signal peptide). Sequences and Uniprot identifiers: RNase T1 (P00651), RNase F1 (P10282), RNase ms (P00653), RNase alpha-sarcin (P00655), Hirsutellin A (P78696), ZT6 (F9X693), SRE1 (R0K2C5), FG12 (I1S329), AVRA1 (N1JGD1), AVRA6 (A0A383UZH9), AVRA7 (N1J8Q9), AVRA9 (N1J9M0), AVRA10 (N1J9C5), AVRA13 (N1JFM8), AVRA22 (N1J9C5), AVRPm2 (A0A1L5JEG4). Amino acid sequence conservation compared to RNase T1 is highlighted in yellow; red Arrows indicate catalytic residues required for ribonuclease activity.

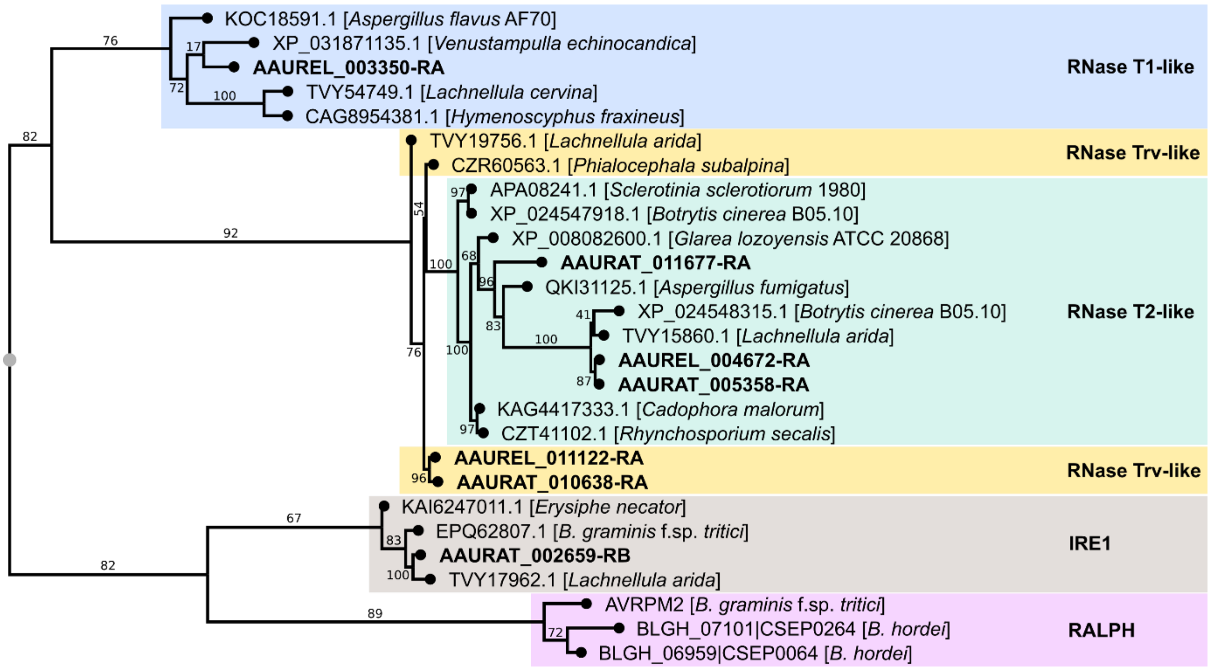


**Supplementary Figure 4. Proteins of *Arachnopeziza aurata* and *A. aurelia* with ribonuclease-like domains are distinct from RNase-like powdery mildew effectors.** *A. aurata* and *A. aurelia* putative secreted proteins containing PFAM domains (PF00445, PF06479) or InterPro domains (IPR036430, IPR016191) from the Ribonuclease T2-like superfamily were queried against the BLAST non-redundant protein database (nr on https://blast.ncbi.nlm.nih.gov/Blast.cgi accessed 04/2025) and protein sequences with similarity extracted for alignment. In addition, three RNase-like effector proteins (AVRPM2 of *B. graminis* f.sp. *tritici* and CSEP0064 and CSEP0264 of *B. hordei* (Spanu, 2017)) were included for multiple sequence alignment. Alignment and phylogenetic reconstructions were performed using the function "build" of ETE3 3.1.3 (Huerta-Cepas *et al.*, 2016) as implemented on the GenomeNet (https://www.genome.jp/tools/ete/; accessed 04/2025). The maximum likelihood (ML) tree was inferred using RAxML v8.2.11 with model PROTGAMMAJTT and default parameters (Stamatakis, 2014). Branch supports were calculated using a Shimodaira-Hasegawa-like approximate likelihood ratio test, a method for assessing the statistical support of different tree topologies (SH-like values). The NCBI GenBank accessions and respective species are indicated; AAUREL indicates *A. aurelia* and AAURAT *A. aurata* proteins. IRE1, Inositol-requiring protein 1 (IRE1); RALPH, RNase-like proteins associated with haustoria; RNase, ribonuclease.

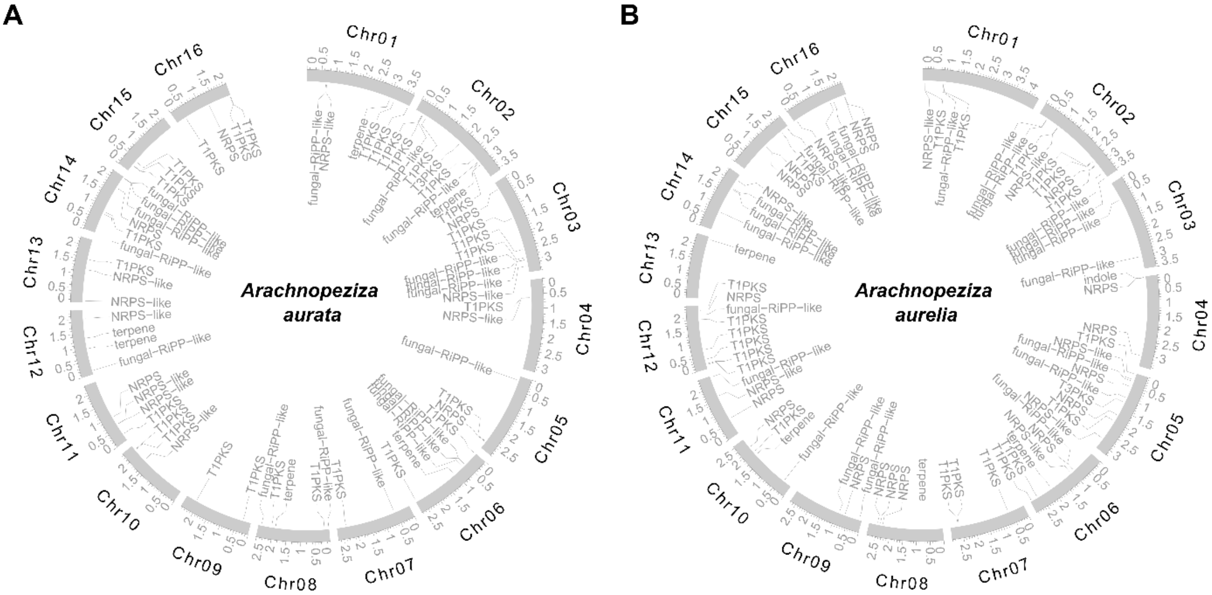


**Supplementary Figure 5. The genomes of *Arachnopeziza aurata* and *A. aurelia* harbor complements of secondary metabolite biosynthesis gene clusters throughout their genomes.** We detected components of secondary metabolism using antiSMASH 7.0 (Medema *et al.*, 2011; Blin *et al.*, 2023) and mapped the respective locations of the coding genes on the genomic maps of *A. aurata* (**A**) and *A. aurelia* (**B**), displayed as circos plots generated with the R package circlize v0.4.10 (Gu *et al.*, 2014). The outer tracks indicate the chromosomal map and the scale the chromosomal position in million base pairs (Mbp). The labels denote the locations of components of secondary metabolism, which were Type 1 polyketide synthase (T1PKS), non-ribosomal peptide synthetase (NRPS, NRPS-like), fungal ribosomally synthesized and post-translationally modified peptide (fungal-RiPP-like), terpene biosynthesis (Terpene), and indole biosynthesis (Indole).

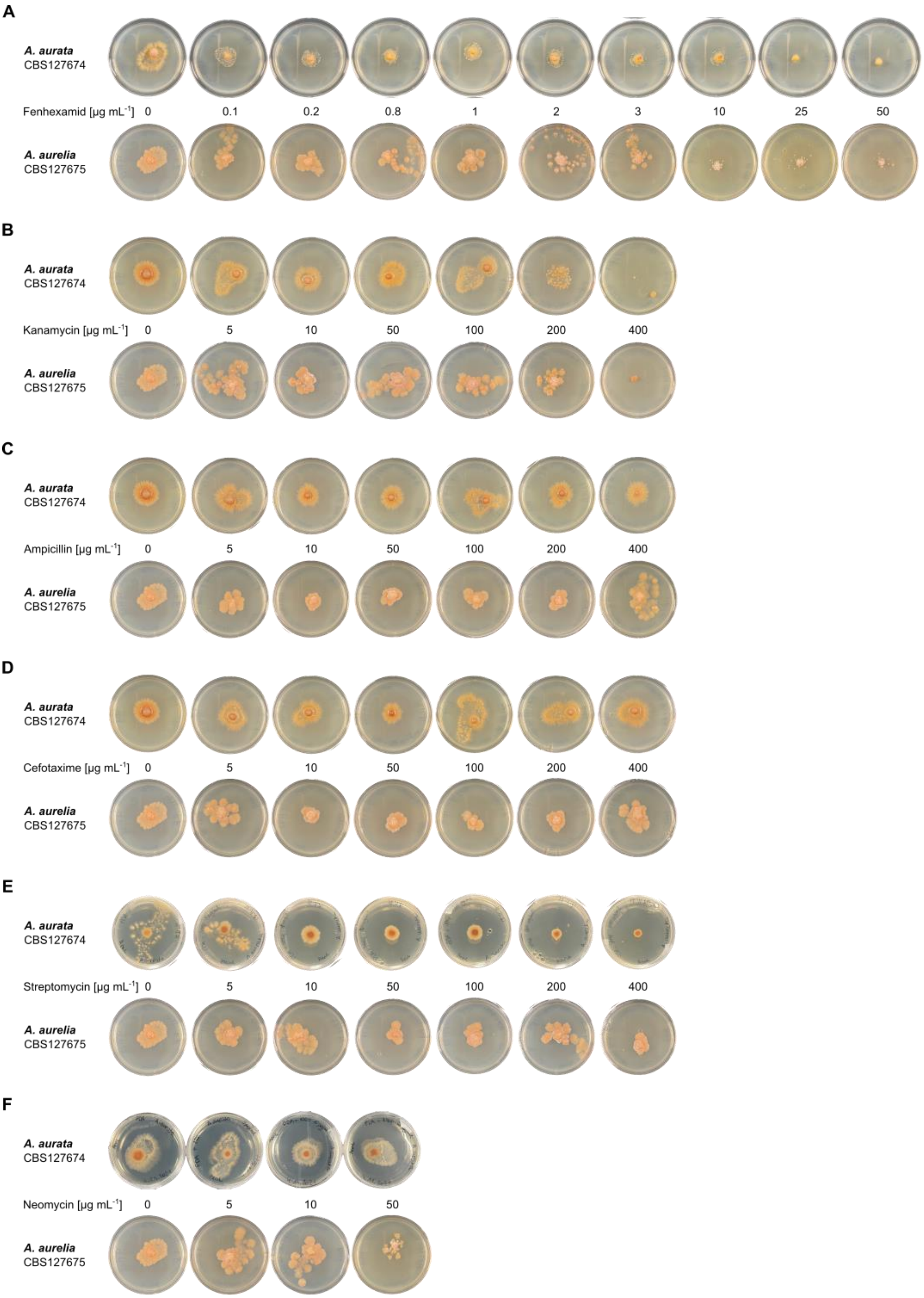


**Supplementary Figure 6. *Arachnopeziza aurata* and *A. aurelia* are inhibited by common fungicides.** The strains *Arachnopeziza aurata* CBS127674 (left) and *A. aurelia* CBS127675 (right) were cultivated on potato dextrose agar (PDA) containing (**A**) fenhexamid, (**B**) kanamycin, (**C**) ampicillin, (**D**) cefotaxime, (**E**) streptomycin, or (**F**) neomycin at the indicated concentrations. The plates were incubated at 23 °C; photographs were taken 16 days after inoculation. Note that the control pictures for the 0 µg $mL^{-1}$ negative control are identical if the respective fungicides/antibiotics were tested in the same experiment.
